# Visual experience drives functional reorganization of cortical GABAergic circuits

**DOI:** 10.64898/2026.02.12.705412

**Authors:** Stefan Sun, Yuxuan Yue, Zimo Li, Quentin Perrenoud, Jessica A. Cardin

**Affiliations:** Department of Neuroscience, Kavli Institute for Neuroscience, Wu Tsai Institute, Yale University School of Medicine, New Haven, CT 06510 USA

## Abstract

Sensory experience shapes the development and function of neocortical circuits. Although repeated exposure to sensory stimuli can alter cortical responses, the precise relationship between sensory experience and circuit interactions remains unclear. Here, we examine how cortical circuits are shaped by repeated exposure to varied visual stimuli. We find that visually evoked beta activity, a prominent pattern of visually evoked activity that relies on somatostatin-expressing (SST) GABAergic interneurons, is robustly potentiated by visual experience. Enhanced visual experience leads to increased responses in SST interneurons and suppressed responses in vasoactive intestinal peptide-expressing (VIP) interneurons. In association with this rebalancing of cortical inhibitory activity, experience increases the recruitment and visual selectivity of excitatory neurons. This enhancement is associated with increased SST-IN influence on nearby pyramidal neurons and requires SST-IN activity during visual experience. Visual experience thus selectively reorganizes adult dendrite-targeting inhibitory circuits, promoting network synchrony and enhancing sensory encoding by excitatory neurons.

## Introduction

Neocortical sensory circuits are crucial for detecting perceptual cues and statistical regularities in the surrounding environment. These circuits exhibit robust experience-driven plasticity that shapes their selectivity at all developmental stages. Visual deprivation during a juvenile critical period induces robust plasticity in the primary visual cortex (V1) that relies on GABAergic inhibition^1–14^. In contrast, loss of sensory input in adulthood leads to cortical plasticity^15,16^ that requires NMDA receptors (NMDARs) but may not rely on GABAergic circuits^15–17^. Enhanced overall sensory experience may also contribute to plasticity in the adult cortex. Indeed, juvenile-like plasticity can be induced in the adult V1 cortex^18,19^ and mice raised in an enriched environment exhibit lifelong plasticity that is lost upon transfer to a standard environment^20–22^. However, how activity in the adult cortex is shaped by visual experience as opposed to deprivation remains poorly understood.

V1 in rodents and other species exhibits distinct learning-related plasticity that can lead to either increased or decreased response magnitudes in excitatory and inhibitory neurons depending on context and reward contingencies^23–36^. Repeated presentation of a single stimulus can evoke a stimulus-specific, NMDAR-dependent potentiation of visually evoked potential (VEP) responses in adult V1^37–39^. This response potentiation relies on GABAergic inhibition and is associated with enhanced thalamocortical input to layer 4 and increased sensory-evoked firing^17,37–44^. In contrast, other work has found that repeated presentation of single stimuli produces robust, NMDAR-dependent loss of response magnitude in excitatory and inhibitory neurons in V1 without changes in thalamocortical inputs^23,45^. The precise relationship between visual experience and adult cortical plasticity thus remains unclear.

In many species, visual stimuli evoke robust rhythmic activity in V1, organizing spiking and enhancing encoding of relevant sensory information^46–58^. Rhythmic synchronization typically arises across distant retinotopic cortical sites when they are stimulated by the same visual patterns^47–49,51,57^, suggesting that the circuits underlying patterned activity are sensitive to the statistics of the sensory environment. Recent work using natural images found that visually evoked rhythms are linked to the predictability of visual features at one retinotopic site compared to the rest of the image^59^. Rhythmic synchrony in V1 may be strengthened when stimuli match an internal representation of expected visual structure that is shaped by sensory experience^60–64^. In mouse visual cortex, the most prominent visually evoked rhythm, beta activity (15-30Hz), relies on dendrite-targeting GABAergic interneurons that co-express the peptide somatostatin (SST-INs)^47,48^ and is regulated by interneurons that co-express vasoactive intestinal peptide (VIP-INs)^49^, suggesting that these inhibitory populations may potentially contribute to experience-dependent reorganization of mature cortical circuits.

Here, we examined the sensitivity of cortical circuits to repeated visual experience of single stimuli or varied stimulus sets by recording V1 local field potentials (LFP) and spiking activity longitudinally over several days. We find that visual experience causes a robust, selective potentiation of visually evoked cortical beta activity, characterized by increased amplitude of individual underlying network events synchronizing activity across cortical layers. Visual experience strengthens the entrainment of cortical neurons by beta events. This form of plasticity is induced by both drifting gratings and natural movies but requires exposure to a rich set of visual stimuli. Using longitudinal calcium imaging of V1 neurons, we find that enriched visual experience transforms GABAergic activity by altering the balance of visually evoked activity in SST- and VIP-INs, leading to increased influence of SST-INs on local pyramidal neurons. In association, visual experience increases the selectivity of local pyramidal neurons (PNs) for visual stimuli. This enhancement of pyramidal neuron responses requires SST-IN activity during visual experience. Together, these data suggest that varied visual experience reorganizes dendrite-targeting interneuron circuits in the adult visual cortex, enhancing network synchrony and visual encoding by excitatory neurons.

## Results

### Visual experience causes plasticity of evoked network events in V1

Previous work has identified both increased^17,37–44^ and decreased^23,24,45^ cortical sensory responses following repeated sensory experience, highlighting a potentially complex relationship between sensory input and plasticity in mature cortical circuits. To gain insight into how sensory experience modulates cortical activity, we first implanted visually naive adult mice with 16-channel silicon probes in V1 and performed longitudinal recordings of the LFP across cortical layers (Figure 1a, Extended Data Figure 1a). Implanted mice underwent a 7-day passive visual experience protocol (Fig. 1b) during which cortical activity was recorded throughout presentation of an extended stimulus set comprised of 64 drifting gratings varying in size, spatial frequency, and contrast (Figure 1a-b). Each session included 960 total stimulus presentations and 80 min of total visual exposure time.

**Figure 1.**
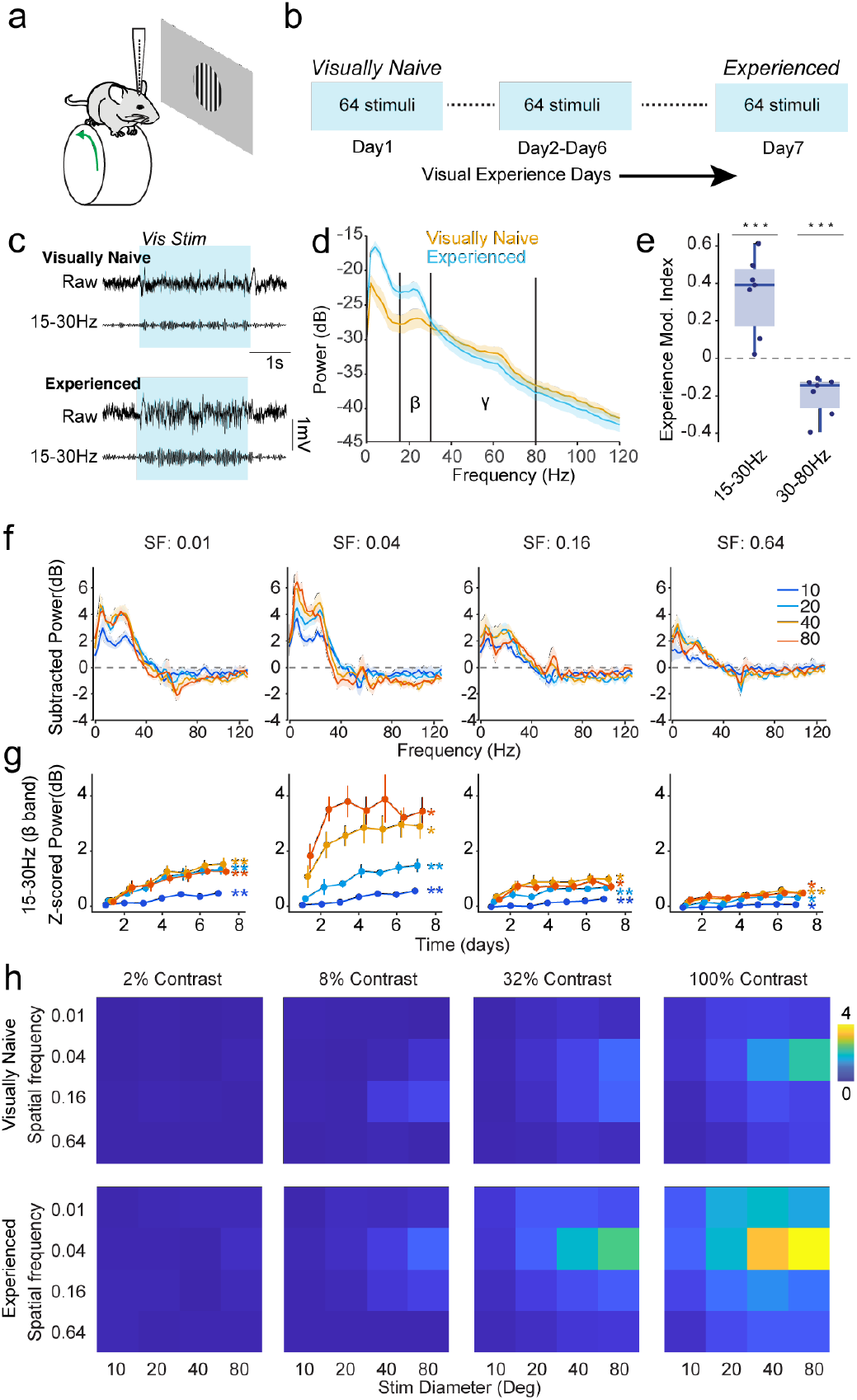
Visual experience potentiates visually evoked beta activity. (a) Experimental paradigm: LFP is recorded across V1 layers with chronically implanted electrode arrays in mice head-fixed on a running wheel while drifting gratings are presented on a screen. (b) Passive visual training schedule: over 7 days, visually naïve mice are exposed to gratings at 4 sizes (10, 20 40 and 80 °), 4 contrasts (2, 8, 32, and 100%) and 4 spatial frequencies (0.01, 0.04, 0,16, 0.64 cycles/°) resulting in a set of 64 stimuli (15 set repetitions; 960 trials per session). (c) Example raw and band-pass filtered (15-30Hz) LFP traces during a large, high-contrast stimulus (80°, 100%, 0.04 cycles/°) in a visually naive mouse (day 1; *upper*) and an experienced mouse (day 7; *lower*). (d) Power spectra of z-scored LFP evoked by a large, high-contrast grating stimulus (80°, 100%, 0.04 cycles/°) in visually naïve (yellow) and experienced (cyan) mice. Experience results in increased power in the beta band (15-30Hz) but not in the gamma band (30-80Hz; n = 7 mice). (e) Experience modulation index for visually evoked power in the beta and gamma bands across all 64 stimulus types (n = 7 mice). (f) Experience-induced change in visually evoked power (Day7 – Day1) for 100% contrast stimuli (16 stimulus types; n = 7 mice). (g) 7-day trajectory of baseline normalized beta power for 100% contrast stimuli (n = 7 mice). (h) Baseline normalized beta power for visually naïve mice (*upper*) and experienced mice (*lower*) across all 64 stimulus types (n = 7 mice). Data are presented as mean ± SEM; *: P<0.05, **: P<0.01,***: P<0.001. See Supplementary Table 1 for detailed statistical analysis. See also Extended Data Figure 1.

Robust activity in the beta range (15-30Hz) is a key hallmark of visually evoked LFP activity in mouse V1^47–50^ and may contribute to long-range synchronization of distant retinotopic areas^49,64^. We observed modest evoked beta activity in response to highly salient stimuli in visually naïve mice. However, visual experience resulted in markedly enhanced visually evoked beta power (Figure 1c, 1d, Extended Data Figure 1b). To quantify the potentiation of activity across stimulus types, we developed an Experience Modulation Index (EMI, see Methods). EMI was positive across stimuli for power in the beta band but not for the adjacent gamma band (30-80Hz; Figure 1e, Extended Data 1f-h), suggesting that visual experience selectively enhances beta activity. Potentiation of visually evoked beta activity was significant during periods of both quiescence and locomotion (Extended Data Figure 1b), suggesting reorganization of underlying circuit interactions rather than enhancement of behavioral state-dependent dynamics. Power in the beta band increased across visual experience days (Figure 1f-1g), exhibiting an immediate increase and plateauing after the second day for stimuli that evoked strong initial beta responses (Extended Data Figure 1g). In contrast, potentiation was more gradual for stimuli that initially evoked weak beta responses.

To further understand the impact of sensory experience on visually evoked activity in V1, we compared overall evoked LFP power, activity in the beta and gamma bands, and visual evoked potential (VEP) amplitudes for each stimulus type across the visual experience paradigm (Extended Data Figure 1c-e). VEPs in V1 layer 4 (L4) arise largely from thalamocortical synaptic input^38^ and increase in magnitude following repeated visual experience with a single stimulus^17,37–43,65,66^. In good agreement with previous studies, we found that the L4 VEPs strengthened over the course of the passive experience protocol (Extended Data Figure 1c-d). In accordance with previous work^47–49^, we found that visually evoked beta activity (15-30Hz) in mouse V1 is maximal for stimuli with a spatial frequency of 0.04 cycles/° and increases with stimulus size and contrast (Figure 1g-h). VEP amplitude was prominent at 0.04 and 0.16 cycles/° spatial frequencies, increased with size and contrast, and also exhibited potentiation across a wide range of stimuli. (Extended Data Figure 1e). In contrast, we observed no change in visually evoked power in the gamma band for any stimulus (Extended Data Figure 1f-h). Overall, these results suggest that visual experience enhances the network interactions underlying cortical beta activity.

### Visual experience selectively potentiates the amplitude of beta events

We recently found that visually evoked beta activity emerges from specific cortical network events occurring rhythmically during visual stimulation^67^. To examine how these events are regulated by visual experience, we employed a recently developed method, CBASS, for reliably detecting network events with high precision in both the frequency and temporal domains^67^ (see Methods). CBASS examines the LFP within a specific frequency band (i.e., beta) and seeks a specific alignment of phase and amplitude across channels that is predictive of a particular state (i.e., visual stimulation). Consistent with our previous findings, the rate of beta events detected by CBASS increased during visual stimulation (Figure 2a, Extended Data Figure 2) and power in the beta range increased during epochs with high beta event rate (Extended Data Figure 2a). The average field around beta events had energy in the beta range (Extended Data Figure 2d) and current source density (CSD) analysis revealed that they were associated with a propagation of activity from layer 4 to layer 2-3 followed by an activation of layers 5 and 6 (Figure 2b).

**Figure 2.**
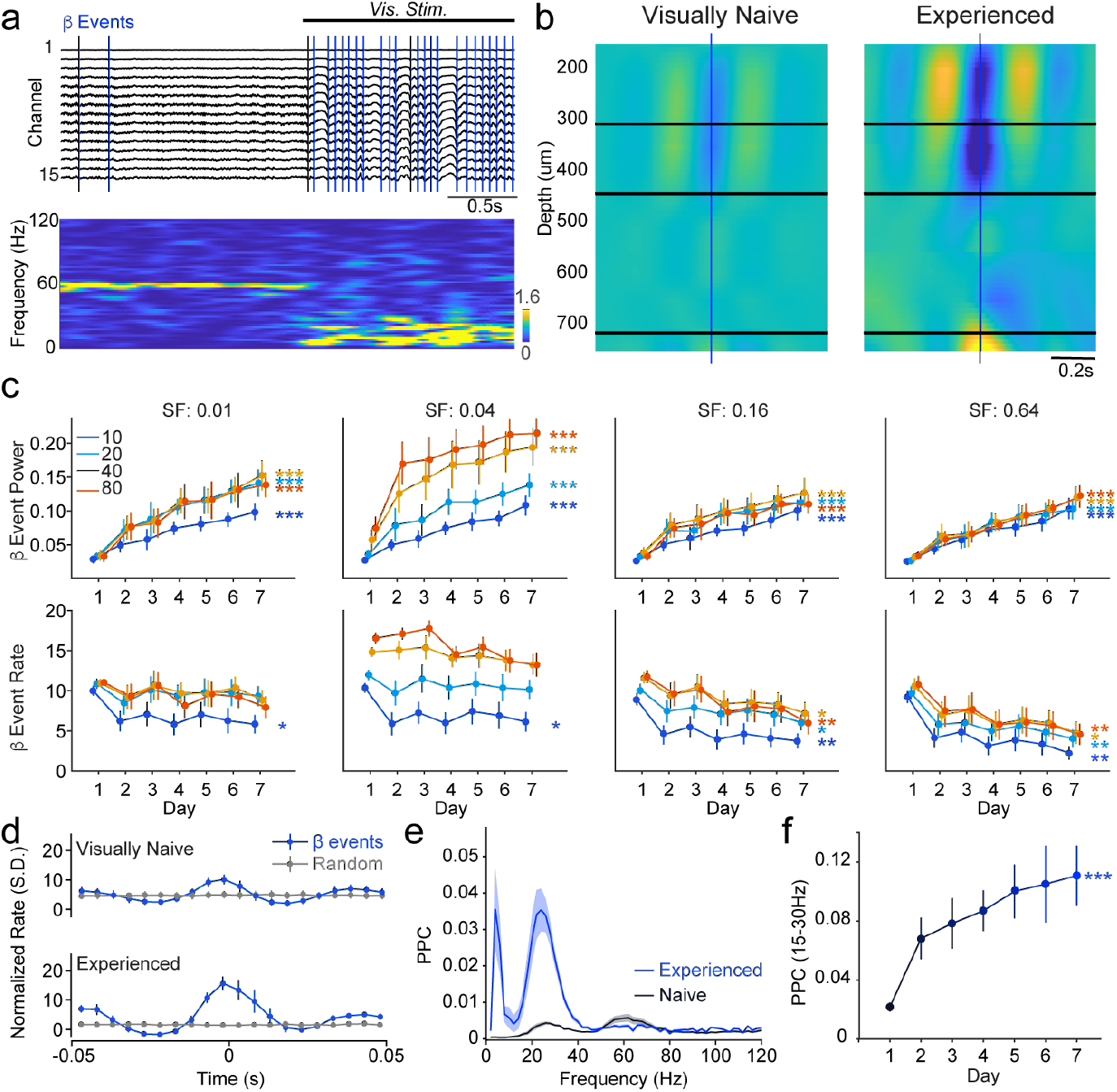
Enhanced beta activity reorganizes visually evoked spiking. (a) LFP across channels (*upper*) and its short time Fourier transform (*lower*) around presentation of a large, high-contrast grating stimulus (80°, 100%, 0.04 cycles/°). CBASS ties visually evoked beta activity to specific network events (blue lines). Note that these events can also happen outside visual stimulation. (b) Averaged CSD around beta events in visually naïve mice (*left*) and experienced mice (*right*; n = 7 mice). (c) 7-day trajectory of beta event power (*upper*) and rate (*lower*) for 100% contrast stimuli at spatial frequencies of 0.01, 0.04, 0.16, and 0.64 cycles per degree (n = 7 mice). (d) Multi-unit activity (MUA) around beta events (blue) or a matched number of random events (gray) in visually naive (*upper*) and experienced mice (*lower*; n = 7 mice). (e) Pairwise Phase Consistency (PPC) spectra quantifying MUA-LFP phase-locking during beta events in visually naive (black) and experienced mice (blue; n = 7 mice). (f) 7-day trajectory of MUA-LFP PPC in the beta range (15-30Hz) during beta events across days of visual exposure (n = 7 mice). Data are presented as mean ± SEM; *: P<0.05, **: P<0.01. See Supplementary Table 1 for detailed statistical analysis. See also Extended Data Figure 2.

Visual experience resulted in a strong potentiation of beta event power (i.e. amplitude squared) across stimulus types (Figure 2b-c) but a slight reduction in beta event rate (Figure 2c). Beta event entrainment of spiking, as measured by multiunit activity (MUA) and spike-LFP phase locking to beta frequencies, increased following visual experience (Figure 2d-f, Methods). These effects were prominent during visual stimulation (Figure 2b-f) and observed during quiescence and locomotion but not in the absence of a visual stimulus (Extended Data Figure 2e-h). Activity in the nearby gamma range can likewise be linked to specific network events detected by CBASS^67^. Gamma event rate increased during locomotion (Extended Data Figure 2i-j) and LFP power increased in the gamma range during epochs with high gamma event rates (Extended Data Figure 2i). Gamma events have faster dynamics, lower amplitude, and weaker activation of layer 6 than beta events (Extended Data Figure 2k). In contrast to beta activity, neither the amplitude nor the rate of gamma events exhibited plasticity in response to the visual experience paradigm (Extended Data Figure 2l). Similarly, visual experience did not alter entrainment of spiking by gamma events (Extended Data Figure 2m-n). Visual experience thus selectively enhances the magnitude of individual visually evoked beta events and the degree to which they organize cortical spiking.

### Experience-dependent plasticity requires exposure to diverse visual stimuli

To examine the conditions necessary to induce experience-dependent potentiation of visual responses, we exposed a cohort of visually naïve mice to repetition of a single highly salient grating stimulus (Extended Data Figure 3a). The full stimulus set was presented on the first and last days of the protocol and only a single stimulus was presented on the intervening days. A second cohort of mice were exposed to a set of 16 full-contrast gratings (Extended Data Figure 3b). Visually evoked beta activity in the 1-stimulus and 16-stimulus cohorts was then compared to that observed in the initial 64-stimulus cohort (Figure 3a). Neither the responses to the repeated single stimulus nor the full stimulus set on the first and last day exhibited plasticity of beta activity in the 1-stimulus cohort (Figure 3a, Extended Data Figure 3a). In turn, we observed an intermediate degree of potentiation in response to the 16-stimulus protocol (Extended Data Figure 3a-b). Potentiation of beta activity was strongest after exposure to the full 64-stimulus protocol (Figure 3b). Likewise, enhancement of beta event power, reduction in beta event rate, and MUA entrainment to beta events were robust following the 64-stimulus protocol, intermediate following the 16-stimulus protocol, and absent following the 1-stimulus protocol (Figure 3c-e). Visual experience-dependent potentiation of evoked beta activity thus requires more than one repeated stimulus and increases with stimulus diversity.

**Figure 3.**
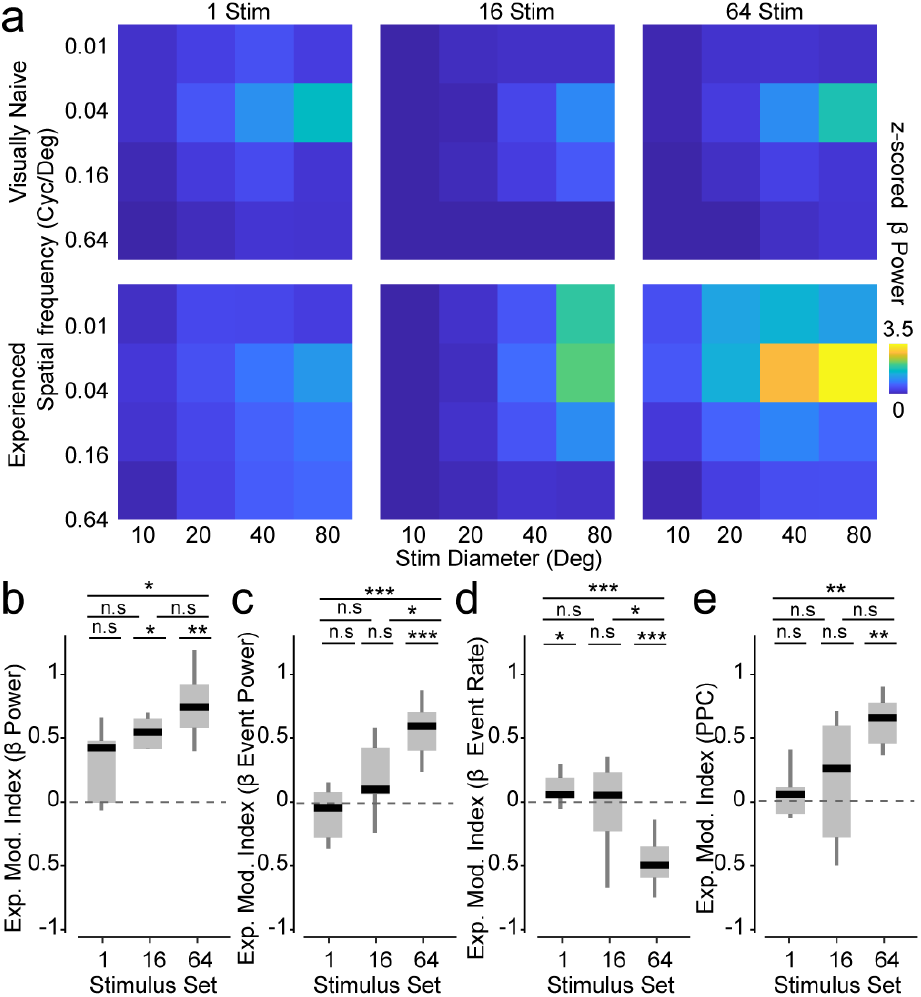
Experience-dependent response potentiation requires diverse stimuli. (a) Baseline normalized beta band power evoked by 100% contrast stimuli in visually naïve mice (day 1; *upper*) and experienced mice (day 7; *bottom*) using 3 different exposure protocols having a matched number of trials (960 trials) between day 2 and 6: *1 Stim.*: repetition of one stimulus (80°, 100%, 0.04 cycles/°; *left*), *16 Stim.*: repetition of all full contrast stimuli (16 stimuli; *middle*) and *64 Stim.*: repetition of the full 64 stimulus sets (*right*; n = 7 mice per protocol). The full stimulus set was displayed on day 1 and day 7 in the 1 Stimulus protocol. (b) Experience modulation index quantifying changes in baseline normalized beta power between day 1 and day 7 across all 100% contrast gratings for each protocol (n = 7 mice per protocol). (c) Same as (b) for beta event power. (d) Same as (b) for beta event rate. (e) Same as (b) for LFP-MUA PPC during beta events. Data are presented as mean ± SEM; *: P<0.05, **: P<0.01, ***: P<0.001. See Supplementary Table 1 for detailed statistical analysis. See also Extended Data Figure 3.

Drifting grating stimuli evoke strong neural responses in V1 across species but represent highly artificial stimulus conditions. We therefore recorded activity in a cohort of visually naïve mice repeatedly exposed to four natural movie clips, each shown at four sizes (Figure 4a-c; see Methods). On the first and last days, mice were also presented with 16 full-contrast gratings. We found that beta activity exhibited plasticity in response to all natural movie clips, and this enhancement was most robust for the largest size stimuli (Figure 4d-e, Extended Data Figure 4a). In contrast, gamma activity in response to the movie clips remained unchanged (Extended Data Figure 4b-c). Beta activity evoked by drifting gratings on the first and last day was also significantly potentiated across all stimuli (Figure 4e). Likewise, the power and entrainment of MUA by beta events increased and their rate decreased after visual exposure in response to both movie clips and grating stimuli (Figure 4f-h). Experience-dependent potentiation of beta activity is thus generic to a broad range of visual statistics and generalizes to stimuli not shown in the repeated set.

**Figure 4.**
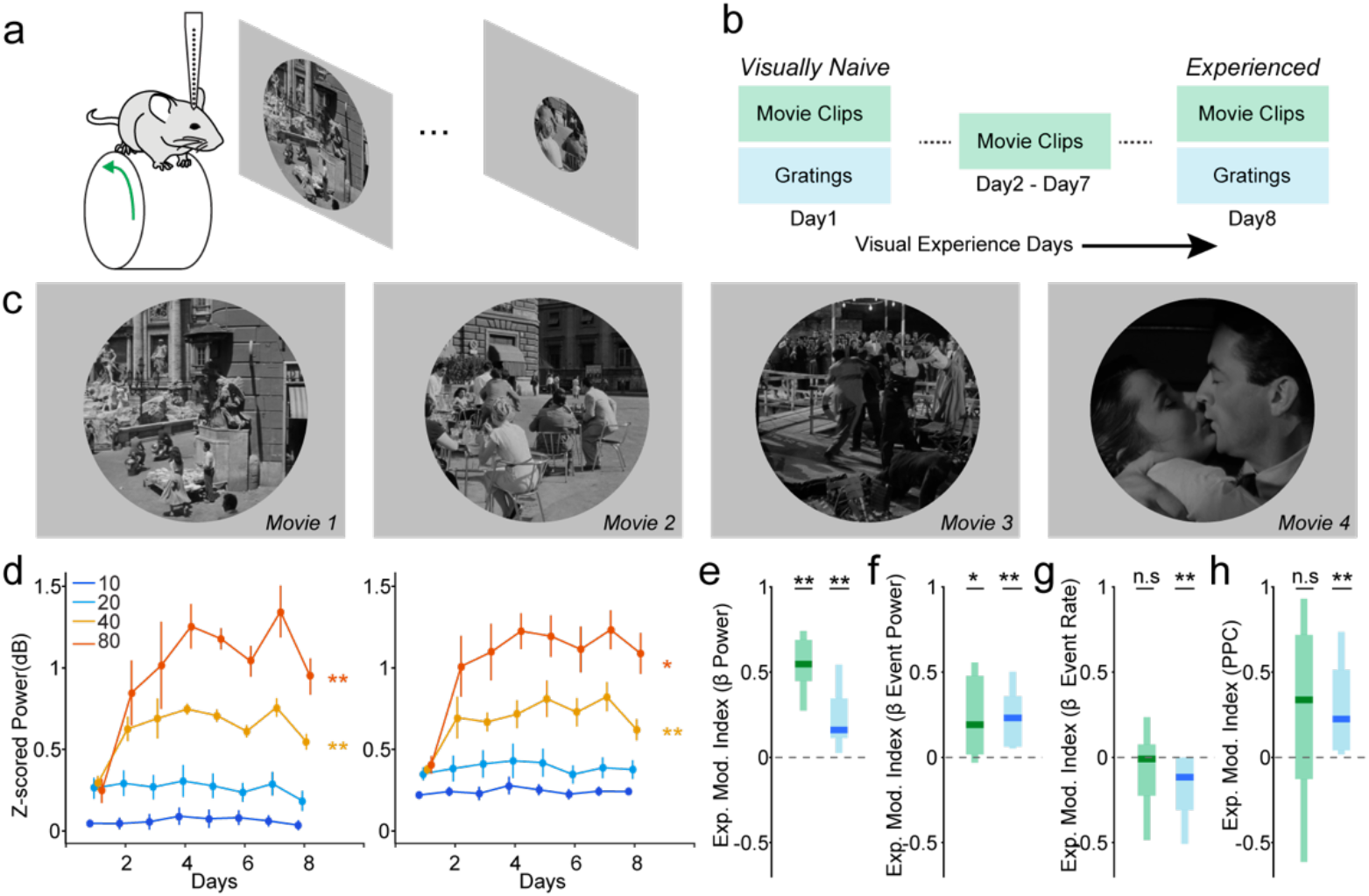
Visually evoked beta activity is potentiated by natural images. (a) Experimental paradigm: LFP is recorded across V1 layers with chronically implanted electrode arrays in head-fixed mice on a running wheel while natural movie clips are presented on a screen. (b) Passive visual training schedule: visually naïve mice are exposed to 4 movie 2s clips at 4 sizes (10, 20, 40, and 80 °) resulting in a set of 16 stimuli (days 1 and 8: 30 set repetitions, 480 trials; days 2 to 7: 60 set repetitions, 960 trials). On days 1 and 8, a set of 16 grating stimuli are also presented at 100% contrast (30 set repetitions; 480 trials). (c) Example frames for each movie clip. (d) 7-day trajectory of baseline normalized beta power evoked by two example movie clips (Movies 3 and 4). Exposure to natural movie clips potentiates beta power. (e) Experience modulation index quantifying changes in baseline normalized beta power between day 1 and day 8 across movie clips (green) and grating stimuli (blue; 7 mice). (f) Same as (e) for beta event power. (g) Same as (e) for beta event rate. (h) Same as (e) for LFP-MUA PPC during beta events. Data are presented as mean ± SEM; *: P<0.05, **: P<0.01, ***: P<0.001. See Supplementary Table 1 for detailed statistical analysis. See also Extended Data Figure 4

### Experience-dependent plasticity of GABAergic interneuron circuits

Previous work has found that visually evoked beta activity in V1 requires the activation of GABAergic interneurons co-expressing somatostatin (SST-INs)^47,48^ and is negatively regulated by interneurons co-expressing vasoactive intestinal peptide (VIP-INs)^49^. SST- and VIP-INs are mutually inhibitory^68–73^, suggesting that expression of beta activity may be dynamically regulated by interactions between these two populations. We therefore examined how visual experience shapes the activity of SST- and VIP-INs in adult V1. We performed 2-photon Ca+ imaging to track visual responses of individual interneurons across days (Figure 5a-b), using the 16-stimulus protocol (Extended Data Figure 3b; see Methods). Neurons were discarded if their receptive fields were misaligned with the visual stimuli (see Methods). Overall visual response amplitudes, pooled across all stimuli, were significantly enhanced by visual experience in approximately half of the retained SST-INs (42/81 cells, 9 mice), whereas smaller proportions exhibited decreases (19/81 cells, 9 mice) or no change (20/81 cells, 9 mice) (Figure 5c-f). The average visual response amplitude across the SST-IN population was robustly enhanced by visual experience (Figure 5g-h). These results remained consistent when analysis was restricted to periods of locomotion (Extended Data Figure 5a-f). Furthermore, SST-INs whose responses were ultimately enhanced or suppressed exhibited distinct trajectories even on the initial days of visual experience and remained separated across days (Extended Data Figure 5m-p), suggesting a rapid and robust differentiation between these subpopulations.

**Figure 5.**
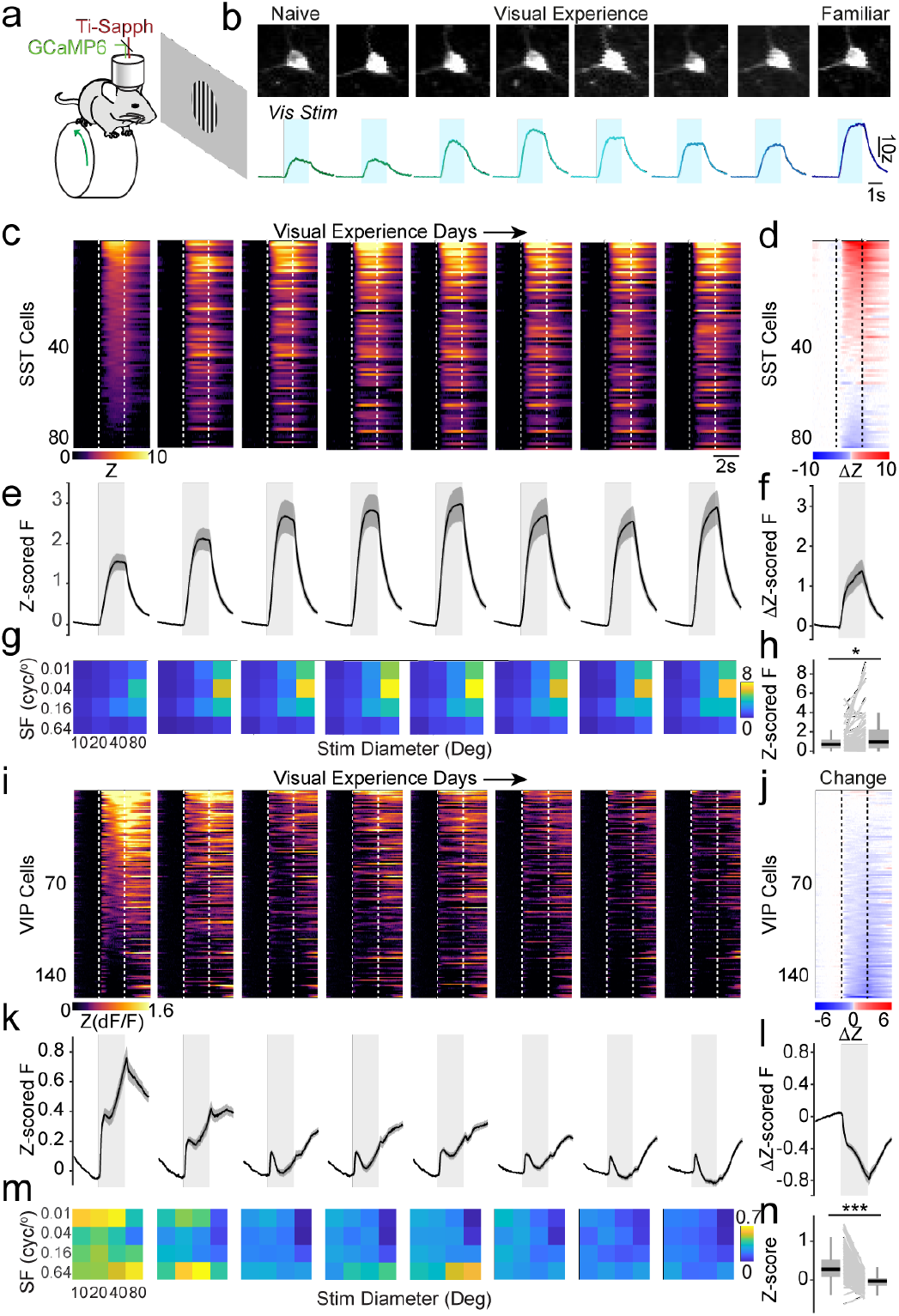
Visual experience induces opposing plasticity regimes in SST and VIP interneurons. (a) Schematic representation of the experimental paradigm. Mice are head-fixed on a running wheel under a 2-photon microscope. Neural responses in V1 layer 2/3 are recorded via a cranial window, during presentation of a series of drifting gratings at 4 sizes (10, 20 40 and 80 °) and 4 spatial frequencies (0.01, 0.04, 0,16, 0.64 cycles/°). (b) Example SST-IN (upper) and its average response to a preferred stimulus (lower) over 8 days of visual experience. Blue shaded area indicates the visual stimulus.(c) Raster heatmap of population calcium activity. Z-scored calcium response averaged across all stimuli, represented by a colormap, around the visual stimulus indicated between dotted vertical lines, in SST-INs for each day of visual experience, sorted by the average response of the cells in naïve mice (first column) (n = 81 cells, from 9 mice). (d) Raster heatmap of the change in calcium response in each cell, calculated by subtracting first-day activity from last-day activity (first and last column in C), sorted by the magnitude of change across days (n = 81 cells, from 9 mice). (e) Z-scored response around visual stimulus averaged across all SST-INs in C, SEM shown in dark grey, visual stimulus window shown in light grey shade (n = 81 cells, from 9 mice). (f) Change in calcium response averaged across all cells in D (n = 81 cells). (g) Average z-scored response across SST interneurons for each combination of size and spatial frequency over days of visual exposure (n = 81 cells). (h) Average z-scored response across visual stimuli for each SST interneurons between naive and visually experienced mice (n = 81 cells). (i) Same as C for VIP interneurons (n = 152 cells). (j) Same as D for VIP interneurons (n = 152 cells). (k) Same as E for VIP interneurons (n = 152 cells). (l) Same as F for VIP interneurons (red: upregulated, n = 8; blue: downregulated, n = 115; black: all, n = 152 cells). (m) Same as G for VIP interneurons (n = 152 cells). (n) Same as H for VIP interneurons (n = 152 cells). Data are presented as mean ± SEM; *: P<0.05, **: P<0.01, ***: P<0.001. See Supplementary Table 1 for detailed statistical analysis. See also Extended Data Figure 5.

In contrast to the SST-INs, the majority of VIP-INs exhibited reduced visual response amplitudes after visual experience (115/152 cells, 7 mice). In a small number of VIP-INs, responses were either enhanced (8/152 cells, 7 mice) or unchanged (29/152 cells, 7 mice) (Figure 5i-l). Average visual response amplitudes across the VIP-IN population were decreased for most visual stimuli following the visual experience protocol (Figure 5m-n). These findings were largely unchanged by locomotion (Extended Data Figure 5g-l). The small subset of VIP interneurons whose responses were enhanced following visual experience were largely those that were suppressed by visual stimulation on the initial days of visual experience (Extended Data Figure 5q-t), suggesting that visual experience causes VIP-INs to reduce their visual responses regardless of the direction of their initial response. In contrast, we found that parvalbumin-expressing interneuron (PV-IN) visual responses were largely unaffected by visual experience (Extended Data Figure 5u-af), showing a small overall decrease in population visual response amplitude that was not replicated during locomotion. Varied visual experience thus has a modest effect on soma-targeting inhibitory circuits. Overall, these results suggest that visual experience primarily rebalances the VIP-SST inhibitory circuit, suppressing VIP-INs and enhancing the visually evoked output of SST-INs.

### Visual experience enhances visual selectivity in pyramidal neurons

SST-INs inhibit the dendrites of PNs, shaping input integration, Ca+ signaling, and synaptic plasticity^69,71,74–76^. To examine how experience-dependent plasticity of SST-IN output affects excitatory neurons in the local circuit, we imaged the activity of layer 2/3 PNs. Visual experience resulted in a redistribution of responses in PNs (Figure 6a-d), with subsets of PNs exhibiting suppression (148/313 cells, 9 mice), enhancement (106/313 cells, 9 mice), or no change (59/313 cells, 9 mice). However, the population average PN visual response amplitude, pooled across all stimuli, remained unchanged (Figure 6e-f). These results remained consistent when analysis was restricted to periods of locomotion (Extended Data Figure 6a-f). PNs whose responses were ultimately enhanced or suppressed began to diverge after the first day of visual experience (Extended Data Figure 6g-j) and exhibited distinct tuning for spatial frequency (Extended Data Figure 6k), suggesting that experience-dependent reorganization of the V1 circuit differentially affects PN populations with different visual selectivity (Extended Data Figure 6k). To examine whether rebalancing GABAergic inhibition also affects the variability of PN responses across stimulus repetitions, we assessed pairwise noise correlations between pyramidal neurons using a noise dependence measure (see Methods). We found that visual experience resulted in a significant decrease in noise dependence across days (Extended Data Figure 6m-n), suggesting that experience increases the efficiency of visual encoding.

**Figure 6.**
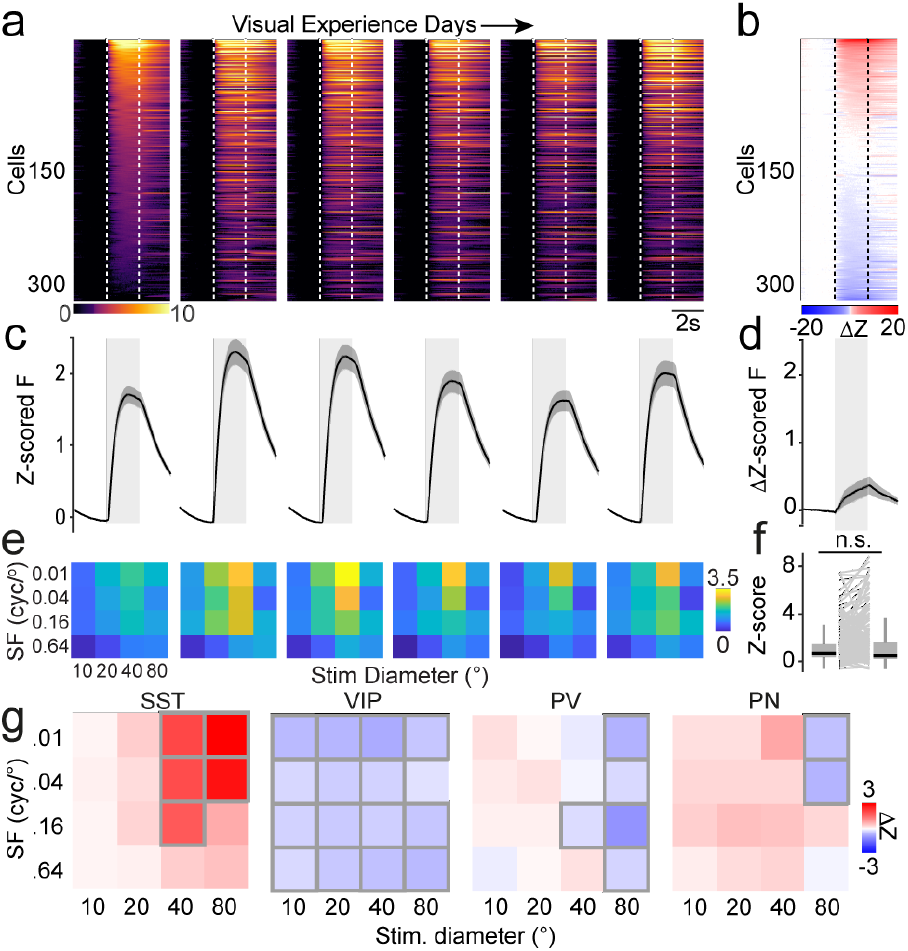
Visual experience enhances responses in pyramidal neurons. (a) Raster images of average z-scored visual response around stimulus onset across stimuli in pyramidal neurons for each day of visual experience, sorted by magnitude of the cell’s initial response in naïve mice (n = 313 cells). (b) Raster image of the average response change around stimulus onset across stimulus type between naive and visually experienced mice in pyramidal neurons, sorted by change magnitude (n = 313 cells). (c) Average z-scored response around stimulus presentation across pyramidal neurons and stimuli over days of visual experience (n = 313 cells). (d) Average change in response between naive and visually experienced mice across all pyramidal neurons (n = 313 cells). (e) Average z-scored response across pyramidal neurons for each combination of size and spatial frequency over days of visual exposure (n = 313 cells). (f) Average z-scored response across visual stimuli for each pyramidal neuron between naive and visually experienced mice (n = 313 cells). (g) Average change in response between naive and visually experienced mice for each combination of size and spatial frequency for SST (n = 81 cells), VIP (n = 152 cells), and PV interneurons (n = 47 cells) and for pyramidal neurons (n = 313 cells). Gray borders denote significance. Data are presented as mean ± SEM; *: P<0.05, **: P<0.01, ***: P<0.001. See Supplementary Table 1 for detailed statistical analysis. See also Extended Data Figure 6.

Previous work found that SST-INs play a key role in mediating surround suppression in pyramidal neurons^77–79^, suggesting that the experience-dependent enhancement of SST-IN output may shape PN selectivity for visual stimuli. To examine how visual experience affects the influence of SST-INs on PNs, we performed a series of experiments combining 2-photon imaging of PNs and targeted optogenetic suppression of nearby SST-INs (Figure 7a-d)^96^. Visual experience in the absence of optogenetic manipulation of SST-INs revealed an increase in PN response amplitudes for small stimuli (≤ 40 degrees) (Figure 7e) and an increase in PN surround suppression index (Figure 7f-g) following the visual experience protocol. Enhanced visual experience with varied stimuli may thus sharpen the selectivity of excitatory neurons for smaller visual stimuli. In a second cohort of animals, we optogenetically suppressed SST-INs during all visual stimulus presentations throughout the visual experience protocol (Figure 7d). Reduction of SST-IN activity during visual stimulation eliminated both the experience-dependent change in PN response amplitude (Figure 7h-i) and the experience-dependent enhancement of surround suppression (Figure 7j). Enhancement of PN selectivity by visual experience thus requires normal visually evoked SST-IN activity. Finally, we used optogenetic suppression of SST-INs on interleaved trials on the first and last days of the paradigm to compare the impact of SST-INs on PN responses before and after visual experience (Figure 7d). Prior to visual experience, suppressing SST-INs selectively enhanced PN responses to large stimuli (Figure 7k). However, after the visual experience paradigm SST-IN suppression also enhanced PN responses to smaller stimuli (Figure 7l-m), suggesting a broader influence of SST inhibition. Together, our results suggest that visual experience reshapes V1 circuit responses and optimizes encoding of stimuli by pyramidal neurons by rebalancing interneuron contributions (Figure 7n).

**Figure 7.**
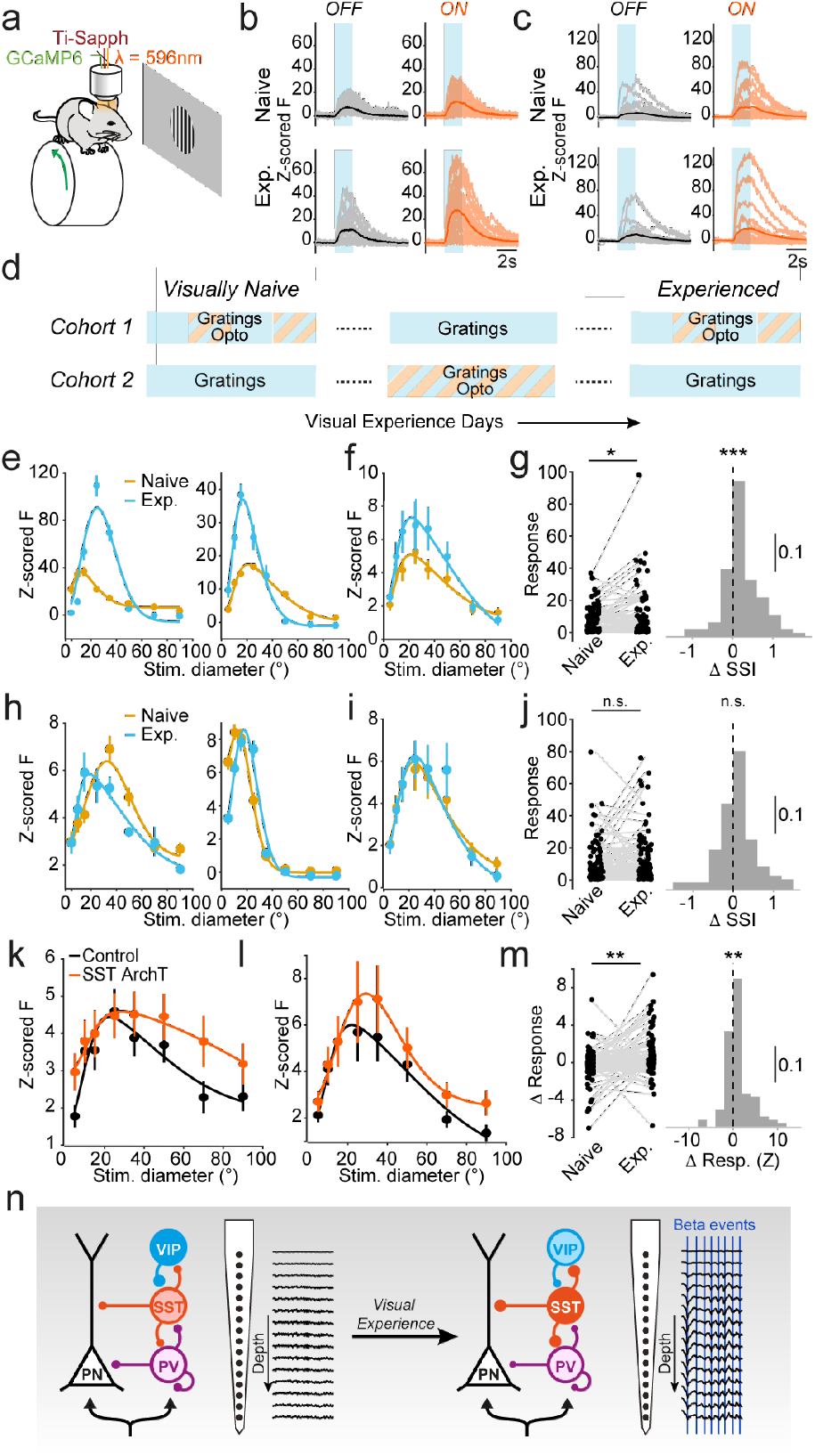
Visual experience sharpens pyramidal neuron selectivity. (a) Schematic of experimental configuration for simultaneous in vivo optogenetics and 2-photon imaging. (b) Responses of an example V1 pyramidal neuron to the same visual stimulus during visually naïve (upper) and visually experienced (lower) periods and without (left) and with (right) optogenetic suppression of SST-INs. (c) Same as b, for a second example pyramidal neuron. (d) Schematic representation of two cohorts of animals in which pyramidal neuron visual responses were assessed during targeted suppression of SST-INs either during a random subset of stimuli only on the first and last days of the paradigm (Cohort 1) or during all visual stimulus presentations except on the first and last days (Cohort 2). (e) Size tuning of two example pyramidal neurons from Cohort 1 in the naive (yellow) and visually experienced (cyan) states. (f) Average size tuning of pyramidal neurons from Cohort 1 in the naive (yellow) and visually experienced (cyan) states (n = 107 cells in 5 mice). (g) Histogram of the change in surround suppression index (SSI) for each pyramidal neuron between the first and last day of the visual experience protocol. (h) Size tuning of two example pyramidal neurons from Cohort 2 in the naive (yellow) and visually experienced (cyan) states. (i) Average size tuning of pyramidal neurons from Cohort 2 in the naive (yellow) and visually experienced (cyan) states (n = 115 cells in 5 mice). (j) Histogram of the change in surround suppression index (SSI) for each Cohort 2 pyramidal neuron between the first and last day of the visual experience protocol. (k) Average size tuning of pyramidal neurons in visually naïve animals from Cohort 1 during visual stimuli without (black) and with (orange) suppression of SST-INs. (l) Same as k, for visually experienced animals from Cohort 1. (m) Histogram of the change in impact of the SST-IN suppression on the amplitude of the pyramidal neuron response to 25° stimuli between visually naïve and visually experienced animals. (n) Schematic model of changes induced in the V1 circuit by varied visual experience. In naive mice (left), SST interneurons are inhibited by visually responsive VIP interneurons, resulting in less SST-IN output to pyramidal neurons and decreased oscillatory synchrony. In visually experienced mice (right), decreased VIP and PV interneuron visually evoked output and potentiated SST output enhances the selectivity of pyramidal neuron visual responses and gives rise to stronger oscillatory synchrony.

## Discussion

Our results reveal a novel form of experience-dependent reorganization in the adult visual cortex that requires exposure to varied stimuli. We find that increased visual experience leads to a robust, selective enhancement of visually evoked beta activity. In response to varied visual stimuli over days, V1 exhibits increased amplitude of individual beta events in the cortical network and increased entrainment of spiking by those events. Underlying this augmentation of visually evoked patterned activity, we find that visual experience induces a reorganization of visual responses in GABAergic interneurons, with SST-IN and VIP-INs exhibiting increased and decreased visually evoked activity, respectively. Visual experience increases the impact of SST-INs on nearby PNs, leading to increases in responses to small stimuli and surround suppression. Together, these changes in circuit interactions lead to enhanced selectivity of excitatory pyramidal neurons. Finally, we find that this enhanced visual selectivity requires SST-IN activity during the visual experience.

Visually evoked rhythmic synchrony in V1 emerges during the display of structured visual stimuli, particularly when visual features at one retinotopic site match that of the surround^47,48,51,52,57,59,64^. Synchrony thus arises when stimuli conform to an internal representation of the expected structure of an image^60–63^. However, it is unclear whether this representation is fixed early in development or remains sensitive to experience in maturity^42^. We find that visually evoked synchrony in the beta range in adult mouse V1 is plastic and strengthened by visual experience. Beta synchrony was induced both by drifting gratings and natural movie clips, suggesting that experience-dependent potentiation does not require a precise match between center and surround but rather a predictable relationship shaped by experience. Repeated exposure to a single static phase-reversing grating is reported to lead to modest potentiation of visually evoked synchrony in the alpha and beta bands in layer 4 V1^42^. However, our results suggest that beta plasticity is strongest upon experience of a varied stimulus set.

Previous work in mouse V1 found that visually evoked beta is reduced by the suppression of SST-INs^42,47,48^ and by activating VIP-INs^49^. Experience-induced potentiation of beta could thus arise from either reduction of VIP-IN activity or enhancement of SST-IN activity. SST-INs are recruited by beta rhythmicity with a delay compared to excitatory and inhibitory fast spiking (FS) neurons^48,50^. Increased SST-IN output may thus strengthen the refractory phase following synchronous discharge during beta events. In accordance with this hypothesis, our data indicate that experience-dependent plasticity increases both SST-IN output and the amplitude of beta events but reduces event rate. In addition, individual beta events more robustly entrained spiking following repeated visual experience, suggesting that beta potentiation reorganizes spike events and enhances spike synchrony in V1.

Repeated presentation of visual stimuli has been shown to result in a range of impacts on the amplitude of visually evoked cortical activity. Presentation of a single phase-reversed grating results in long-lasting potentiation of the amplitude of the layer 4 VEP that emerges across sessions, along with increased visually evoked firing of layer 4 excitatory neurons^37–39,65,80,81^. This potentiation can be evoked by a range of stimuli but does not generalize to novel stimuli^37,38^ and is associated with enhanced power in broadband alpha and beta frequencies (8-30Hz) and reduced gamma range activity^24,36,42^. In contrast, repeated passive presentation of a single drifting grating stimulus results in decreased visually evoked responses in layer 4 and layer 2/3 excitatory neurons across days^23,24,45,65^, suggesting that moving stimuli may lead to adaptation rather than potentiation of visual responses. Here, we found that repeated presentation of single and varied drifting grating stimuli and natural movies all resulted in robust potentiation of layer 4 VEPs. Because thalamocortical synaptic inputs make a substantial contribution to layer 4 VEPs, these results suggest that repeated presentation of a wide range of stimuli may potentiate the amplitude or synchrony of thalamocortical inputs. However, potentiation of visually evoked activity patterns within the cortex may instead depend on the number of stimulus repetitions or variation in stimulus features.

Variation in wakeful behavioral state, such as between quiescence and locomotion, regulates the amplitude of visual responses^82–85^ and plays a role in experience-dependent plasticity^33–35^. In addition, repeated stress can alter cortical sensory responses^86,87^. Earlier studies incorporated up to two days of habituation to restraint prior to beginning repeated visual stimulation sessions^17,37–42,65^ and recorded cortical activity in mice restrained in a tube^2,6,15,17,19,37–43,65,66,80,81^. In contrast, our protocol included extensive habituation and recordings in mice able to freely run on a wheel. Differences in either baseline behavioral state or locomotion may thus account for some reported differences across paradigms. However, we found that the response potentiation induced by repeated visual experience was robust to behavioral state, suggesting that this enhancement does not rely on arousal signals.

Cellular imaging revealed that in response to repeated exposure to varied stimuli, GABAergic interneuron circuits in V1 exhibited substantial changes in visually evoked activity. VIP- and SST-INs represent two distinct populations of GABAergic interneurons in the cortex, with distinct inputs, firing properties, and synaptic targets^68,88–93^. VIP-INs primarily synapse on other GABAergic cells, including SST-INs, and make some synapses directly on PNs^68,69,71,73,94^. In turn, SST-INs inhibit VIP-INs and innervate the dendrites of PNs, contributing to synaptic integration, calcium signaling and plasticity, and potentially gating inputs to apical dendritic segments^68,74–76^. Both VIP- and SST-INs are sensitive to visual input, but whereas VIP-INs exhibit relatively small receptive fields^95^, SST-INs are selective for larger stimuli^77–79,96^. We found that a majority of V1 SST-INs exhibited progressive increases in visual response amplitude across visual presentation sessions. In contrast, almost all VIP-INs exhibited a decrease in responses over the same time period. The soma-targeting PV-INs, which receive robust innervation from SST-INs^68,97^ and may receive input from VIP-INs^72,98^, exhibited a modest decrease in visual responses following visual experience. In contrast, previous work found that repeated presentation of single stimuli leads to progressively decreased activation of layer 4 and layer 2/3 PV interneurons^23,42,45,99,100^. Indeed, stimulus-specific response potentiation to phase-reversed grating stimuli may rely on activation of NMDA receptors in PV-INs^17^ and Nrst1 expressing cortico-thalamic neurons in layer 6^40,41,101,102^. Together, these results suggest that varied visual experience may have a unique impact on visual cortical circuits, selectively reshaping the dendrite-targeting VIP-SST circuit and leading to enhanced SST-IN output to local PN dendrites.

Following repeated experience with varied stimuli, the responses of V1 PNs exhibited a selective increase in responses to small stimuli and increased surround suppression. Surround suppression in V1 has been suggested to emerge in part via inhibition from SST-INs, whose larger receptive fields may give rise to suppression of weaker PN responses to large diameter stimuli while permitting expression of strong PN responses to small diameter stimuli^77–79,96^. Optogenetic suppression revealed that varied visual experience increases the impact of SST-INs on PN visual responses. Targeted suppression of SST-INs during the visual experience paradigm eliminated the experience-dependent enhancement of PN visual responses, suggesting that visually evoked SST-IN activity is critical for the induction of changes in PN visual selectivity. Increased SST-IN output to PN dendrites following visual experience may thus lead to increased surround suppression of PNs and enhanced PN tuning for stimulus size. We further found that visual experience led to decreased pairwise noise dependence between local PNs, suggesting increased efficiency of visual encoding^103,104^. Indeed, previous computational modeling results indicate that increased GABAergic inhibition may reduce noise correlations^105^.

Taken together, our results establish that experience leads to a sustained reshaping of the mature cortical response to visual stimuli. Exposure to varied stimuli leads to rebalancing of the evoked activity in GABAergic IN circuits, enhancing the visually evoked output from SST-INs. These circuit-level changes in inhibitory activation give rise to a robust strengthening of visually evoked beta rhythmicity and reorganization of the pattern of visually evoked spikes. In parallel, enhanced dendrite-targeting output from SST-INs is also associated with enhanced surround suppression and increased visual selectivity in excitatory neurons. This remodeling can be induced by a wide range of stimulus paradigms including drifting gratings and natural stimuli. These findings strongly suggest that the full expression of adult visual experience-dependent plasticity is contingent on specific aspects of the stimulus paradigm such as the statistical properties of the visual stimuli, stimulus repetition, and behavioral state. Future studies should aim to further explore how the precise parameters of sensory stimulation shape circuit interactions in the mature cortex and how different plasticity mechanisms engaged by sensory input may functionally interact to shape visual processing.

## Methods

### Experiments

#### Animals and viruses

All animal procedures were approved by the Institutional Animal Care and Use Committee of the Yale University School of Medicine (Protocol # 11317). Male and female of the following lines were used: C57Bl/6 mice, Sst-IRES-Cre mice (Jax 018973) crossed with Ai148 ((TIT2L-GC6f-ICL-tTA2)-D; Jax 030328), Vip-IRES-Cre mice (Jax 031628) crossed with Ai148, Pvalb-IRES-Cre mice (Jax 017320) crossed with Ai148, Thy1-GCaMP6s mice (Jax 024275), Emx1-cre mice (Jax 075934) injected with AAV9-Syn-FLEX-GCaMP6s (2 × 10^12^ gc ml^−1^; Addgene #100845) and Sst-IRES-Cre^+/-^ injected with AAV9-CaMKII-GCaMP6s (2 × 10^12^ gc ml^−1^; Addgene #107790) and AAV9-CAG-Flex-ArchT (2 × 10^12^ gc ml^−1^; Addgene #209779). Mice were maintained on a 12-h light/dark cycle with ad libitum access to food and water. All experiments were conducted during the light phase when animals were between 3 and 6 months of age.

#### Surgery

Mice were anesthetized with isoflurane (1.5% in oxygen) and maintained at 37°C for the duration of the surgery. Analgesia was provided with subcutaneous injections of Carprofen (5mg/kg). Lidocaine (1% in 0.9% NaCl) was injected under the scalp to provide topical analgesia. Eyes were protected from desiccation with ointment (Puralube). The scalp was resected, and the skull cleaned with Betadine. A surgical screw was implanted on the skull between the eyes and nuts were glued to the skull above the bregma suture, allowing the fixation of a headplate with bolts. Alternatively, the headplate were directly fixed onto the skull with C & B Metabond, (Butler Schein).

For chronic electrophysiology, two craniotomies were performed respectively above V1 on the left hemisphere (∼0.15mm diameter; 3.75mm posterior, 2.5mm lateral from Bregma) and above the cerebellum (0.4mm diameter; ∼6mm posterior from Bregma). An A16 probe with a CM16 connector (Neuronexus) was lowered into V1. Ground and reference wires were inserted above the cerebellum. Craniotomies were protected with Gelfoam (Pfizer), and all implants were affixed to the skull with C & B Metabond, (Butler Schein).

For calcium imaging, a 3 mm^2^ square craniotomy was made over V1 on the left hemisphere. A custom-made glass window consisting of a 3 mm^2^ square inner cover slip and 5 mm^2^ round outer cover slip (both #1, Warner Instruments) glued with an ultraviolet-curing adhesive (Norland Products) was inserted into the craniotomy and secured to the skull with Cyanoacrylate glue (Loctite). A circular ring was glued to the titanium headpost and Metabond applied to cover all exposed skull.

For local virus injections, virus was first loaded into a glass micropipette. The tip of the pipette was lowered into the primary visual cortex through a craniotomy (3.75mm posterior, 2.5mm lateral from Bregma) at a depth of ∼350 μm (QSI, Stoelting Co.).

Analgesics (5 mg/kg Carprofen) and anti-inflammatory steroids (2 mg/mL Dexamethasone) were given immediately after surgery and on the two following days and implanted mice were housed with food containing antibiotics (Sulfatrim, Butler Schein).

#### Chronic in-vivo extracellular electrophysiology

Mice were habituated to handling and head fixation for 5 days prior to recordings. Mice were head-fixed and freely running on a cylindrical wheel equipped with a magnetic angle sensor (Digikey) to monitor wheel motion. The face (including the pupil and whiskers) was imaged with a miniature CMOS camera (Blackfly s-USB3, Flir) with a frame rate of 10 Hz.

For chronic electrophysiology, implants were connected to the recording apparatus (DigitalLynx system, Neuralynx). The most superficial contact point was used as a reference. Local Field Potentials, wheel motion, and timing signals for face movies and visual stimulus were acquired at a 40KHz sampling rate.

#### Chronic in-vivo two-photon cellular imaging

Imaging was performed using a resonant scanner-based two-photon microscope (MOM, Sutter Instruments) coupled to a Ti:Sapphire laser (MaiTai DeepSee, Spectra Physics) tuned to 920 nm for GCaMP6. Emitted light was collected using a 25 × 1.05 NA objective (Olympus). To prevent light contamination, the objective was enclosed in blackout material extending to the headpost. Images were acquired using ScanImage 4.2 at 30 Hz, 512 × 512 pixels at 150–250 μm depth from pia matter. Visual stimulation, wheel position, and Ca2+ imaging microscope resonant scanner frame ticks were digitized (5 kHz) and collected through a Power 1401 (CED) acquisition board using the Spike 2 software.

#### Visual stimulation hardware

Visual stimuli were generated using the Psychtoolbox Matlab extension and displayed on a 17’’ by 9.5’’ monitor situated 15 cm from the right eye. Screen display was linearized, and maximum luminance was adjusted to ∼140 cd.sr/m^2^. An iso-luminant grey background was displayed between visual stimuli.

#### Visual stimulation protocol

Visual exposure protocols lasted 6-8 days and used either vertical gratings drifting leftward at 1 or 2Hz or natural movie clips cropped from the movie *Roman Holiday* (1953). Stimuli were centered on the retinotopic location evoking the strongest visual response, measured with either MUA or visually evoked potential in LFP for electrophysiology, or fluorescence signal for calcium imaging. The following protocols were used:

-64 Stimuli: Gratings were presented for 3s with a 2s inter-stimulus-interval (ITI) at all 64 combinations of 4 sizes (10, 20, 40, 80 ° diameter), contrasts (2, 8, 32 and 100%) and spatial frequencies (0.01, 0.04, 0.16 and 0.64 cycle/°). Each stimulus was repeated 15 times resulting in 960 total trials. This protocol was repeated 7 days.

-1 Stimulus: A single highly salient grating (80°, 100%, 0.04 cycle/°), was presented for 3s with a 2s ITI for 960 trials between days 2 and 6. The 64-stimuli set was displayed on day 1 and 7 to probe for change across grating types.

-16 Stimuli: Gratings were presented for 2s with a 5s ITI at full contrast at all 16 combinations of 4 sizes (10, 20, 40, 80 ° diameter) and spatial frequencies (0.01, 0.04, 0.16 and 0.64 cycle/°). Each stimulus was repeated 60 times resulting in 960 total trials. This protocol was repeated 7 days for electrophysiology recordings and 8 days for 2P recordings. This protocol was selected for 2P calcium recordings over the 64 stimuli since neuronal calcium responses were strongly modulated by behavioral states, thus requiring more repetitions for each stimulus.

-Natural Movies: 4 movie clips lasting 2s were presented with a 5s ITI, masked at 4 sizes (10, 20, 40, 80 ° diameter) resulting in 16 stimulus types. Between days 2 and 7 each stimulus was repeated 60 times resulting in 960 total trials. On days 1 and 7 each stimulus was repeated 30 times and 30 repetitions of the 16 Stimuli set were also displayed to test for changes across grating types, resulting in 960 total trials.

#### Optogenetic stimulation

Optogenetic stimulation was achieved by aligning a 594nm laser to the same light path as the Ti:Sapphire laser of the two-photon microscope, allowing us to activate ArchT without affecting imaging quality. The main dichroic of the microscope was replaced with one with a 594nm notch to allow dual IR and 594nm excitation. Optogenetic stimulation was delivered through the two-photon microscope objective lens as a single pulse of light beginning 200ms prior to the onset of the visual stimulus and concluding 200ms after the end of the 2-second visual stimulus period. To avoid optical noise from fluorophores at higher stimulation levels, an untagged AAV-CAG-ArchT construct was used^96^. The power of 594nm laser was measured at 20mW out of the objective.

## Data Analysis

### Wheel and whisker motion

The first principal component of nose and whisker pad motion energy was computed from each movie of the mouse face using FaceMap^106^. Epochs of running and whisking were defined using a change point algorithm detecting local changes in the mean and variance of running speed and whisker pad motion. Briefly, the moving standard deviation of running speed or the first principal component of whisker motion were computed within a defined temporal window. The length *t* of this window determines the temporal resolution of the change-point analysis and was set to 4s for running speed and 500ms for facial motion. A first estimate of locomotion motion onset/offset times were then taken as the time when the moving standard deviations exceeded/ fell below 20% of its range above minimum. Estimates were refined in a window *t* around each onset/offset time by computing the time points corresponding to the maximum of the *t*-windowed moving forward/backward Z-score.

### In-vivo electrophysiology

Data were analyzed in Matlab 2020b (Mathworks) using custom scripts. All time-series were down-sampled to 1KHz and aligned. Local field potential (LFP) recordings were high-pass filtered at 1Hz using a 2nd order Bessel filter. Multi-unit activity (MUA) was extracted from LFP recording using spikedetekt and klustakwik2 and the phygui (https://github.com/cortex-lab/phy) to extract unsorted spikes.

### Layer alignment of LFP and current source density (CSD) across recordings

To compute the average field potential around CBASS events across recordings, the LFP was linearly interpolated across channels to a common grid of laminar position. The CSD was derived as the second spatial derivative of the LFP across interpolated laminar positions. LFP channels were then mapped onto cortical layers using the CSD profile of visual responses. Two current sinks were identified and assigned respectively to layer 4 and layer 5b^67^.

### Visually Evoked Potential (VEP)

The VEP is defined as the peak to trough amplitude of the LFP around the visual stimulus onset. Specifically, the trough was restricted within 0-100 ms from the stimulus onset, while the peak was restricted within 50-150 ms from the stimulus onset.

### CBASS

CBASS (Clustering Band-limited Activity by State and Spectral features) ties a power increase in a defined frequency band (i.e., gamma (30-80Hz)) during a particular state (i.e., running) to the occurrence of defined events in the temporal domain (ref). A detailed description and implementations in matlab and python are available on (https://github.com/cardin-higley-lab/CBASS). Briefly, the multichannel LFP is filtered in the band of interest and candidate events are selected at the troughs of the filtered signal in a reference channel. In our case, the reference channel was chosen to be closest to layer 4. The spectrotemporal dynamics underlying each candidate event are parameterized using the real and imaginary part of the analytical representation (matlab function *hilbert*) of the filtered signal in each channel. Candidate events form a cloud in this parametric space where neighbors have similar spectro-temporal dynamics The event cloud is split randomly into ***n*** partitions and a binomial test is performed in each partition to determine if events happen during the state of interest (i.e. running) at higher frequencies than overall. Partitioning is repeated ***N*** time. A state enrichment score is calculated for events as the fraction of time they fell into an enriched partition. An optimization procedure is then applied to find the threshold yielding the most significant distance between events having a low and a high enrichment score in the feature space. Events above threshold are retained. Here we used ***n*** = 20 partitions and ***N*** = 1000. Different settings for these parameters have only a marginal influence on the result of the procedure.

To check the validity of the event partition, a state enrichment score is computed as described above on surrogate data having a spectrum and channel covariance matched with that of each recording. This surrogate data is constructed by decomposing the original LFP recording into principal components across channels (matlab function *pca*), randomizing the phase of their Fourier transform and remixing them. The fraction of candidate events above threshold on surrogate data indicates how likely pattern may be associated to the state of interest (i.e. locomotion) given the statistics of the signal.

### Comparison of network activity within and outside CBASS event cycles

CBASS events are aligned to the trough of the band-pass filtered LFP in a reference channel. We defined each event’s boundaries as the peaks surrounding the event’s trough. Peak and troughs were determined as the 0 and π valued time points of the argument (matlab function *abs*) of the analytic representation (matlab function *hilbert*). Activity inside the event boundaries thus fell within a cycle centered on the trough. Epochs during and outside all CBASS event cycles were pooled separately and compared.

### MUA distribution around CBASS events

MUA distribution around CBASS events was computed as follows. The lag separating each MUA spike from the nearest CBASS events was estimated. A histogram of lag values was then computed and normalized as follows. A baseline MUA rate ***p*** was computed over samples. The variance of the rate over a window of ***n*** samples was estimated assuming a binomial distribution as 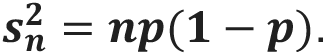 The normalized rate of events over a window of ***n*** samples was then taken as 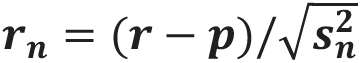 where ***r*** is the event rate over samples and can be thought of as the number of standard deviations away from baseline. Thus, normalized histogram values represent standard deviation away from the mean MUA rate under the assumption of an unmodulated binomial process.

### Spectral analysis

The spectral power of a given time series was derived with Welch’s method. Each channel was divided into 500ms overlapping segments (75% overlap). Each segment was multiplied by a Hamming window and their Fourier transform was computed (matlab function fft). Power was derived as 10 times log10 of the squared magnitude of the Fourier Transform and expressed in dB. Power was averaged over segment and channels.

Band-limited LFP power was estimated by applying the Hilbert transform to zero-phase, band-pass filtered signals and computing instantaneous power as the squared magnitude of the analytic signal. Power was averaged across channels and stimulus epochs. Baseline power statistics were obtained from quiescent, no-stimulus periods, and stimulus evoked power was normalized by z-scoring relative to the baseline mean and standard deviation.

The spectral power of event-triggered averages was derived with a minimum bias multi-taper estimate. This differs from a classical multi-taper estimate in that Slepian tapers are replaced by a sinusoidal tapers sequence defined as:

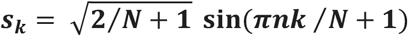

where ***N*** is the number of samples in the triggered average, ***n*** is the sample number and ***k*** is the order of the taper. Sinusoidal tapers produce a spectral concentration almost comparable to that achieved with a Slepian sequence while markedly reducing local bias. The number of tapers was chosen to yield a bandwidth of 8Hz following the formula: ***K*** = ***round***((**4*πNB*** / ***r***) − **1**) where ***B*** is the bandwidth and ***r*** is the sample rate. Triggered averages were multiplied by each taper. Spectral power was then computed as described above and averaged over tapers.

For MUA phase locking quantification a spectro-temporal representations was first derived either for a set of frequencies using a wavelet transform (matlab function *cwt*) and a Morlet wavelet (matlab identifier *cmor1-2*) or across a full frequency band by computing the analytical representation of the filtered signal (matlab function *Hilbert*).

Spike phase locking was estimated using the Pairwise Phase Consistency defined as:

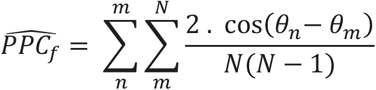

where ***θ_k_*** is the phase of the signal for frequency ***f*** at the time of spike ***k*** and ***N*** is the total number of spikes. PPC provides an unbiased estimate of spike phase locking. However, the estimate can be noisy if the spike number is inferior to 250. Thus, population estimates of PPC were derived by pooling MUA from all selected channels and the variance across mice was estimated by first calculating the PPC for individual mice.

### Calcium signal pre-processing

Analysis of imaging data was performed either using ImageJ and custom routines in MATLAB (Mathworks) or with Suite2p^107^. In the first case, motion artifacts and drifts in Ca2+ signals were corrected with the moco plug-in in ImageJ^108^, and regions of interest (ROIs) were selected as previously described^109^. Pixels in each ROI were averaged and the neuropil signal was subtracted^109–111^. We observed no differences between the two pre-processing pipelines.

### Calcium response to visual stimulation

The visual response was computed as the *z*-scored change in fluorescence during the 2 s visual stimulus (*F*) compared to the 1 s baseline before the stimulus (*F*_0_), given by (*F*-mean(*F*_0)_)/std(*F*_0)_.

### Visual responsiveness

The averaged z-score during the pre-stimulus 1-second period and 2-second stimulus period from all the trials were examined with a paired t-test to determine if each neuron was responsive to each stimulus on each session/day. Neurons that were responsive to a stimulus for more than a day-threshold were defined as responsive to that stimulus. Neurons responsive to at least one 20-degree or smaller size of stimulus, regardless of the spatial frequency, were defined as center-responsive. Only center-responsive neurons were included in the results.

### Locomotion/Quiescent trials for cellular calcium recording

Wheel position was determined from the output of the linear angle detector. The circular wheel position variable was first transformed to the [-π, π] interval. The phases were then circularly unwrapped to get running distance as a linear variable, and locomotion speed was computed as the derivative of distance (cm/s). A change-point detection algorithm detected locomotion onset/offset times based on changes in standard deviation of speed. Locomotion onset or offset times were estimated from periods when the moving standard deviations, as determined in a 0.5s window, exceeded or fell below an empirical threshold of 0.1. Transition points were removed if a locomotion bout is under 1 second or a quiescent bout is under 2 seconds. Locomotion bouts were required to have average speed exceeding 0.5 cm/s. Only trials that completely fall into a locomotion bout or quiescent bout are categorized as a locomotion trial or quiescent trial respectively.

### EMI

Exposure Modulation Index was calculated using Z-scored Beta/Gamma power or calcium fluorescence with equation below:

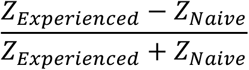

### Noise Correlation and Noise Dependence of Neuron Pairs

For each pair of neurons *i*, *j*, the stimulus-specific noise correlation coefficient *μ_i_*_,j,*s*_ was defined as the Pearson correlation coefficient between their trial-averaged z-scored responses for stimuli *s*:

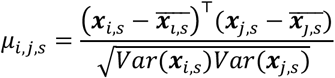

where ***x_i_***_,***s***_ = {***x_i_***_,***s***,**1**_, ⋯, ***x_i_***_,***s***,***T***_} ∈ ***R^T^*** refers to the average z-scored response of neuron ***i*** to stimulus ***s*** in each of the ***T*** quiescent trails. The noise dependence ***d_i_***_,***j***_ is the absolute value of the stimulus-specific noise correlation coefficient, averaged across all stimuli (S=16), such that it only captures the magnitude of shared trial-to-trial variability, independent of the correlation sign:

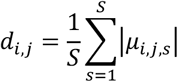

### Statistics

A detailed description of the statistics test in each figure panel and of statistical samples is provided in Extended Data Table 1. Unless otherwise noted, permutation tests were performed using mice as the statistical unit. Multiple comparisons were corrected using the Benjamini & Hochberg procedure for false discovery rate.

### Mixed-effects models

To assess changes in visual responses while accounting for the nested structure of the data (neurons recorded from the same mouse), we used linear mixed-effects (LME) models. First, to test whether the visual response change was significantly different from zero irrespective of stimulus identity, we fit an LME model with the change in response (deltaZ) as the dependent variable, an intercept as the fixed effect, and mouse identity included as a random intercept (deltaZ ∼ 1 + (1 | MouseID)).

Second, to evaluate stimulus-specific response changes, we fit a separate LME model in which stimulus identity was included as a fixed effect without an intercept, allowing direct estimation of the mean response change for each stimulus, with mouse identity again included as a random intercept to account for repeated measurements within animals (deltaZ ∼-1 + Stimulus + (1 | MouseID)). Statistical significance of response changes was assessed using the estimated fixed-effect coefficients from the fitted models.

## Acknowledgements

The authors thank all members of the Higley and Cardin laboratories for helpful input throughout all stages of this study. We thank the Yale Vision Core’s viral core (R.Pant, M. Higley, J. Demb) for generation of AAV vectors and Rima Pant for histology. This work was supported by funding from the NIH (R01EY022951, R01EY035127, and R01MH113852 to JAC, EY026878 to the Yale Vision Core).

## Author Contributions

SS, QP, and JAC designed the experiments. SS, YY and QP collected the data and SS, ZL, and QP analyzed the data. SS, QP, and JAC wrote the manuscript.

## Data Availability Statement

The full datasets generated and analyzed in this study are available from the corresponding authors on request.

## Code Availability Statement

Custom written MATLAB scripts used in this study are available: https://github.com/cardin-higley-lab/Sun-et-al-2026, https://doi.org/10.5281/zenodo.22031129.

---

**Extended Data Figure 1.**
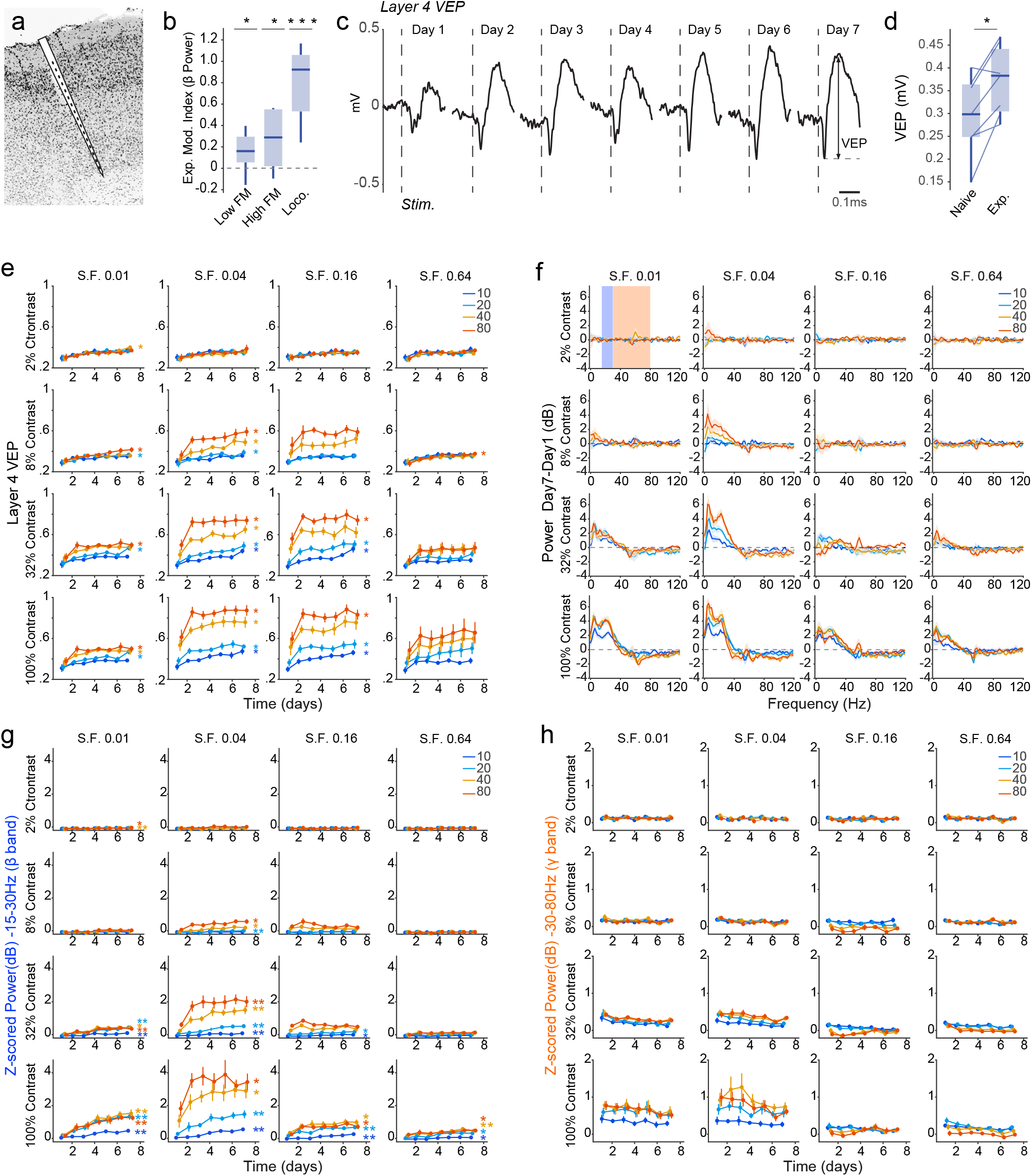
Experience-dependent potentiation of evoked responses across cortical layers. (a) Inverted contrast image of a DAPI stained histological section and schematic showing the location of the multichannel silicon probe in V1 in an example brain. (b) Experience modulation index for visually evoked power in the beta band across all 64 stimulus types during epochs of low facial motion (Low FM), high facial motion (High FM) and locomotion (Loco.) (n = 7 mice). (c) Average layer 4 voltage around visual stimulus onset for highly salient stimuli (80°, 100%, 0.04 cycles/°) in an example mouse across days of visual exposure (n = 15 trials). Visually Evoked Potential (VEP) is defined as the peak-to-trough amplitude within 200ms after stimulus onset. (d) Layer 4 VEP across all 64 stimulus types on day 1 and day 7 (n = 7 mice). (e) 7-day trajectory of layer 4 VEP power for all 64 stimulus types (n = 7 mice). (f) Experience-induced change in visually evoked power (Day7 – Day1) for all 64 stimulus types (n = 7 mice). (g) 7-day trajectory of baseline normalized beta power (15-30Hz) for all 64 stimulus types (n = 7 mice). Note that data for 100% contrast stimuli are replicated in Figure 1 g. (h) 7-day trajectory of baseline normalized gamma power (30-80Hz) for all 64 stimulus types (n = 7 mice). Data are presented as mean ± SEM; *: P<0.05, **: P<0.01, ***: P<0.001. See Supplementary Table 1 for detailed statistical analysis.

**Extended Data Figure 2.**
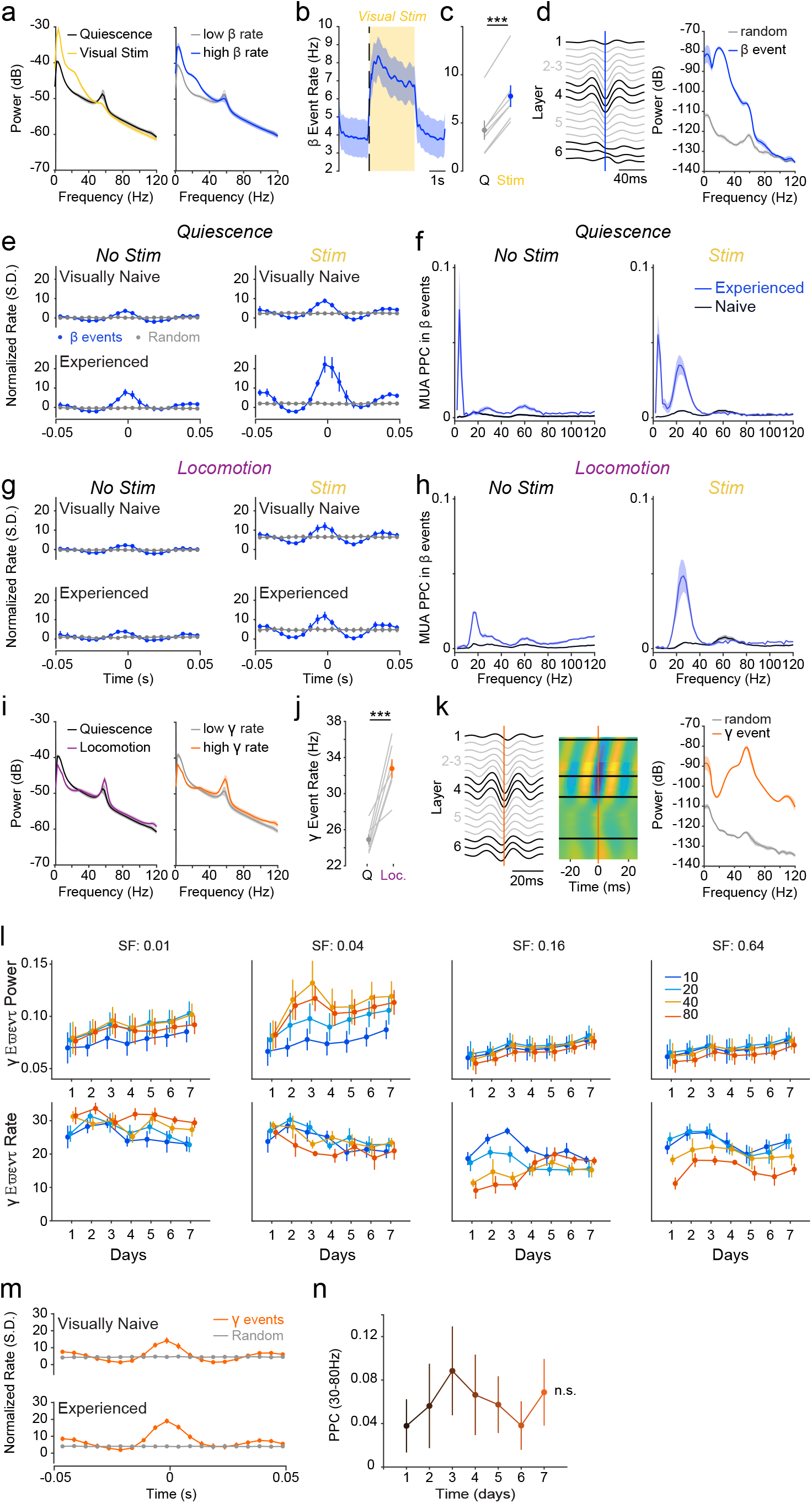
Enhanced beta activity is tied to the potentiation of specific translaminar activity motifs. (a) LFP power spectrum outside visual stimulation (black) and during large, high-contrast stimuli (yellow 80°, 100%, 0.04 cycles/°) (*left*) and during epochs of low (gray) and high (blue) beta event rate (*right*; n = 7 mice). (b) β Event rate around presentation of highly salient stimuli (80°, 100%, 0.04 cycles/°, n = 7 mice). (c) β event rate increases during presentation of highly salient stimuli (80°, 100%, 0.04 cycles/°, n = 7 mice). (d) Average LFP around β events (*left*) and its power spectrum (blue) compared to that of a matched number of random events (gray; *right*; n = 7 mice). (e) MUA around beta events (blue) or a matched number of random events (gray) in visually naive (*upper*) and experienced mice (*lower*) in the absence of visual stimulus (*left*) and during salient visual stimuli (*right*; contrast >=32%) during quiescence (n = 7 mice). (f) Pairwise Phase Consistency (PPC) spectra quantifying MUA-LFP phase-locking during beta events in visually naive (black) and experienced mice (blue) in the absence of visual stimulus (*left*) and during strong visual stimuli (*right*; contrast >=32%) during quiescence (n = 7 mice). (g) Same as E during locomotion. (h) Same as F during locomotion. (i) LFP power spectrum during quiescence (black) and locomotion (purple; *left*) and during epochs of low (gray) and high (orange) gamma event rate (*right*; n = 7 mice). (j) Gamma event rate increases during locomotion (n = 7 mice). (k) Average LFP around gamma events (*left*), associated CSD (*middle*) and power spectrum of the average field (orange) compared to that of a matched number of random event (gray; *right*; n = 7 mice). (l) 7-day trajectory of gamma events power (*upper*) and rate (*lower*) for 100% contrast stimuli (n = 7 mice). (m) MUA around gamma events (orange) or a matched number of random events (gray) in visually naive (*upper*) and experienced mice (*lower*; n = 7 mice). (n) 7-day trajectory of MUA-LFP PPC in the gamma range (30-80Hz) during gamma events across days of visual exposure (n = 7 mice). Data are presented as mean ± SEM; ***: P<0.001; black bar: significant after correction for multiple comparison. See Supplementary Table 1 for detailed statistical analysis.

**Extended Data Figure 3.**
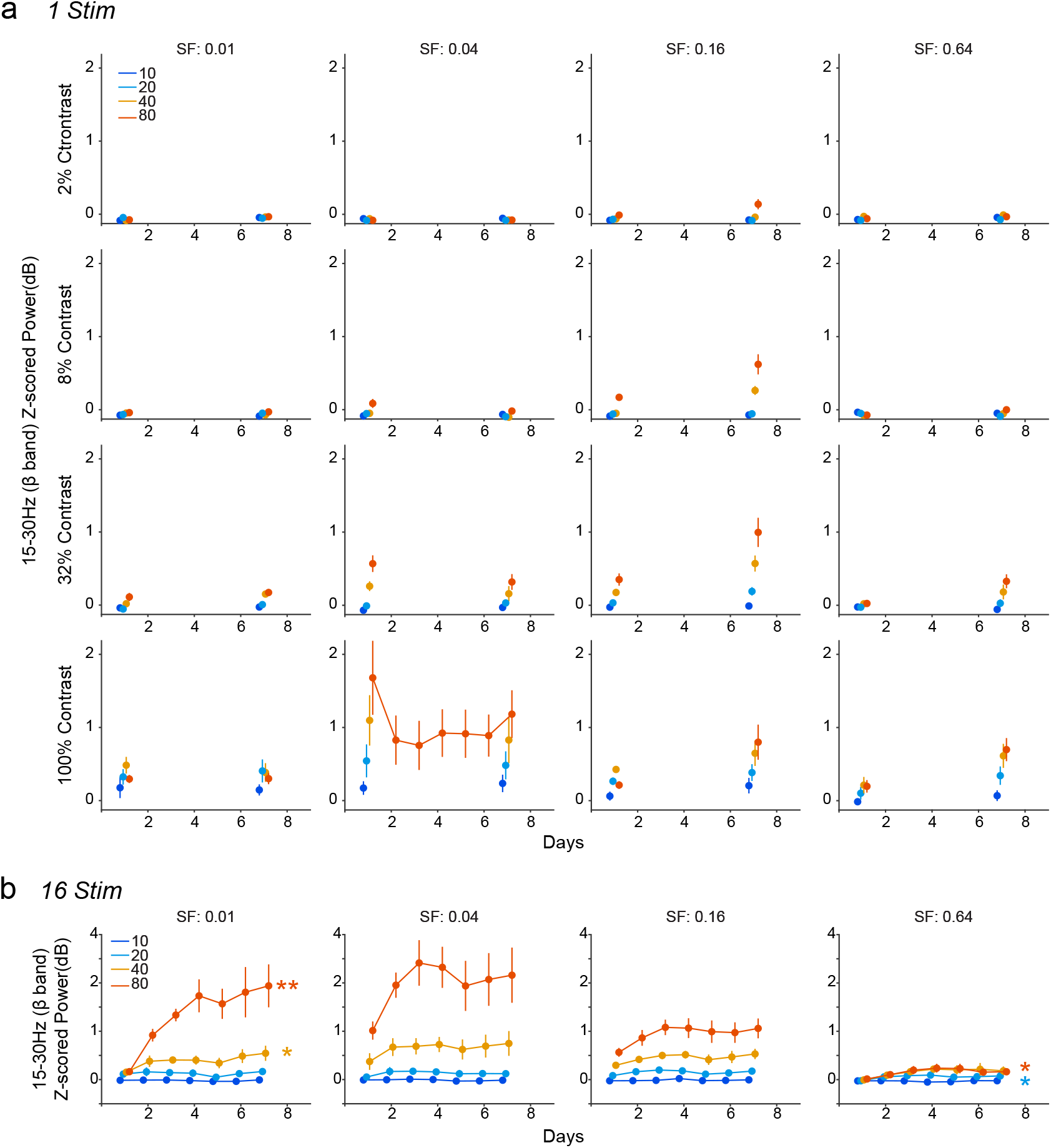
Detailed trajectories of cortical response to each stimulus paradigm. (a) 7-day trajectory of baseline normalized beta power (15-30Hz) for all 64 stimulus types in the 1 Stimulus visual exposure protocol (n = 7 mice). (b) 7-day trajectory of baseline normalized beta power (15-30Hz) for all 16 stimulus types in the 16 Stimulus visual exposure protocol (n = 7 mice). Data are presented as mean ± SEM; *: P<0.05, **. See Supplementary Table 1 for detailed statistical analysis.

**Extended Data Figure 4.**
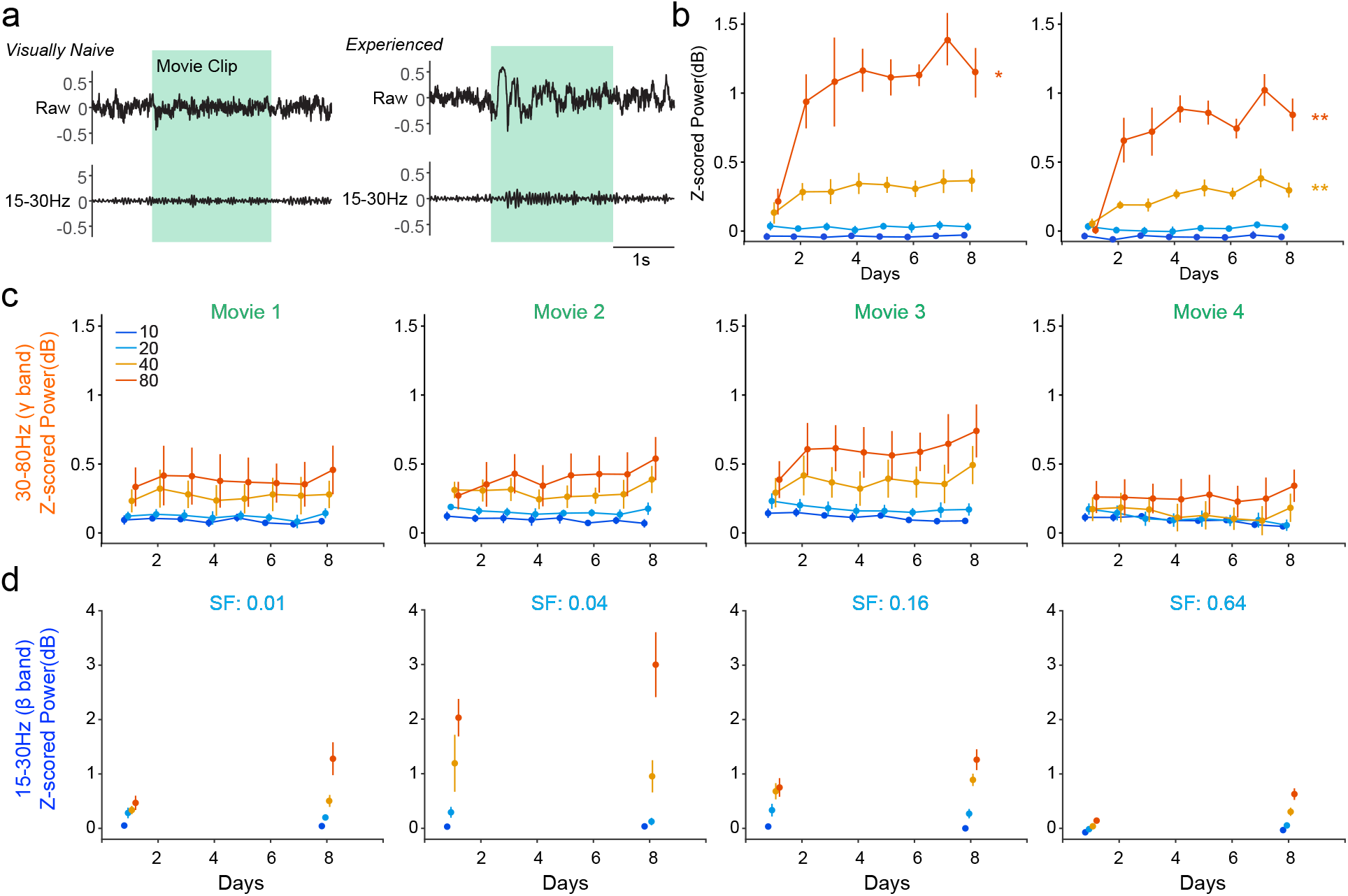
Visually evoked beta activity is potentiated by natural images. (a) Example raw and band-pass filtered (15-30Hz) LFP traces during a natural movie clip in a visually naive mouse (day 1; *left*) and an experienced mouse (day 7; *right*). (b) 7-day trajectory of baseline normalized beta power evoked by two example movie clips (Movies 1 and 2). (c) 7-day trajectory of baseline normalized gamma power evoked by each movie clip. Exposure to natural movie clips does not significantly potentiate gamma power (n = 7 mice). (d) 7-day trajectory of baseline normalized beta power evoked by all 16 grating stimuli used at day 1 and day 8. Exposure to movie clips does not significantly potentiate beta power during the display of grating stimuli (n = 7 mice). Data are presented as mean ± SEM. See Supplementary Table 1 for detailed statistical analysis.

**Extended Data Figure 5.**
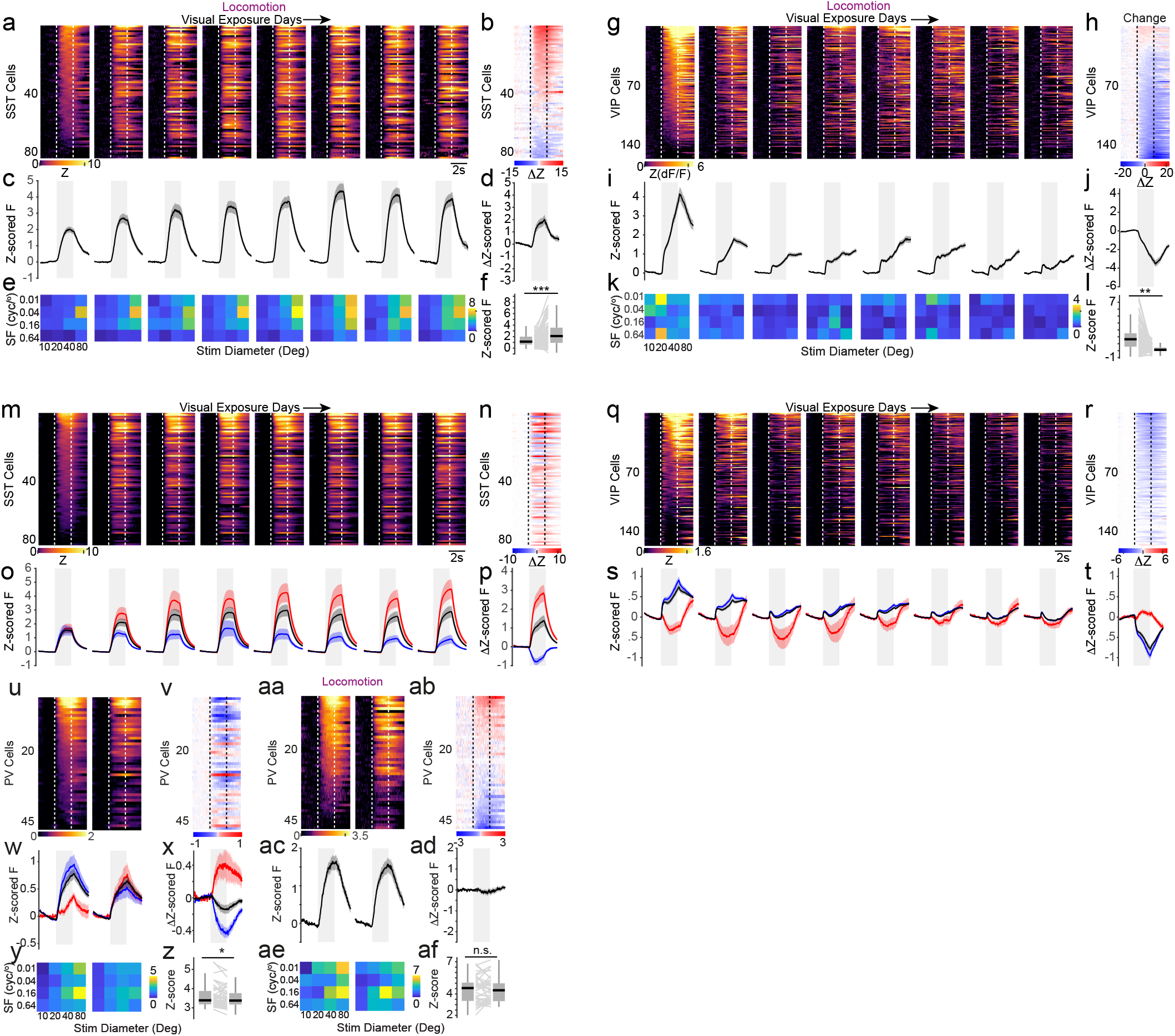
Visual experience leads to distinct changes in VIP- and SST-IN visual response profiles. (a) Raster images of average z-scored response around stimulus onset across stimulus in SST interneurons for each day of visual exposure sorted by magnitude of response in naïve mice, restricting analysis to periods of locomotion (n = 81 cells). (b) Raster image of the average response change around stimulus onset across stimulus type between naïve and visually experienced mice in SST interneurons sorted by change magnitude, restricting analysis to periods of locomotion (n = 81 cells). (c) Average z-scored response around stimulus presentation across SST interneurons and stimuli over days of visual experience, restricting analysis to periods of locomotion (n = 81 cells). (d) Average change in response between naive and visually experienced mice in upregulated (red, n = 42 cells) and down regulated SST interneurons (blue, n = 19 cells) and across all SST interneurons (black, n = 81 cells), restricting analysis to locomotion. (e) Average z-scored response across SST interneurons for each combination of size and spatial frequency over days of visual exposure (n = 81 cells), restricting analysis to locomotion. (f) Average z-scored response across visual stimuli for each SST interneurons between naive and visually experienced mice (n = 81 cells), restricting analysis to locomotion. (g) Same as a, for VIP interneurons (n = 152 cells). (h) Same as b, for VIP interneurons (n = 152 cells). (i) Same as c, for VIP interneurons (n = 152 cells). (j) Same as d, for VIP interneurons (red: upregulated, n = 8; blue: downregulated, n = 115; black: all, n = 152 cells). (k) Same as e, for VIP interneurons (n = 152 cells). (l) Same as f, for VIP interneurons (n = 152 cells). (m) Raster images of average z-scored response around stimulus onset across stimulus in SST cells for each day of visual exposure, sorted by change magnitude (n = 81 cells). (n) Raster image of the average response change around stimulus onset across stimulus type between naïve and visually experienced mice in SST cells, sorted by change magnitude (n = 81 cells). (o) Average z-scored response around stimulus presentation across stimuli over days of visual exposure in upregulated (red, n = 42 cells) and downregulated SST interneurons (blue, n = 19 cells) and across all SST interneurons (black, n = 81 cells). (p) Average change in response between naive and visually experienced mice in upregulated (red, n = 42 cells) and down regulated SST interneurons (blue, n = 19 cells) and across all SST interneurons (black, n = 81 cells). (q) Same as m, for VIP interneurons (n = 152 cells). (r) Same as n, for VIP interneurons (n = 152 cells). (s) Same as o, for VIP interneurons (red: upregulated, n = 8; blue: downregulated, n = 115; black: all, n = 152 cells). (t) Same as p, for VIP interneurons (red: upregulated, n = 8; blue: downregulated, n = 115; black: all, n = 152 cells). (u) Raster images of average z-scored response around stimulus onset across stimulus in PV interneurons for the first and last day of visual exposure, sorted by magnitude of response in naïve mice (n = 47 cells). (v) Raster image of the average response change around stimulus onset across stimulus type between naïve and visually experienced mice in PV interneurons, sorted by change magnitude (n = 47 cells). (w) Average z-scored response around stimulus presentation across stimuli on the first and last days of visual exposure in upregulated (red, n = 5 cells) and downregulated PV interneurons (blue, n = 16 cells) and across all PV interneurons (n = 47 cells). (x) Average change in response between naive and visually experienced mice in upregulated (red, n = 5 cells) and down regulated PV interneurons (blue, n = 16 cells) and across all PV interneurons (black, n = 47 cells). (y) Average z-scored response across PV interneurons for each combination of size and spatial frequency on the first and last days of visual exposure (n = 47 cells). (z) Average z-scored response across visual stimuli for each PV interneurons between naive and visually experienced mice (n = 47 cells). (aa) Same as u, restricting analysis to locomotion (n = 47 cells). (ab) Same as v, restricting analysis to locomotion (n = 47 cells). (ac) Average z-scored response around stimulus presentation across stimuli on the first and last days of visual exposure, restricting analysis to locomotion (n = 47 cells). (ad) Average change in response between naive and visually experienced mice, restricting analysis to locomotion (n = 47 cells). (ae) Same as y, restricting analysis to locomotion (n = 47 cells). (af) Same as z, restricting analysis to locomotion (n = 47 cells). Data are presented as mean ± SEM; **: P<0.01, ***: P<0.001. See Supplementary Table 1 for detailed statistical analysis.

**Extended Data Figure 6.**
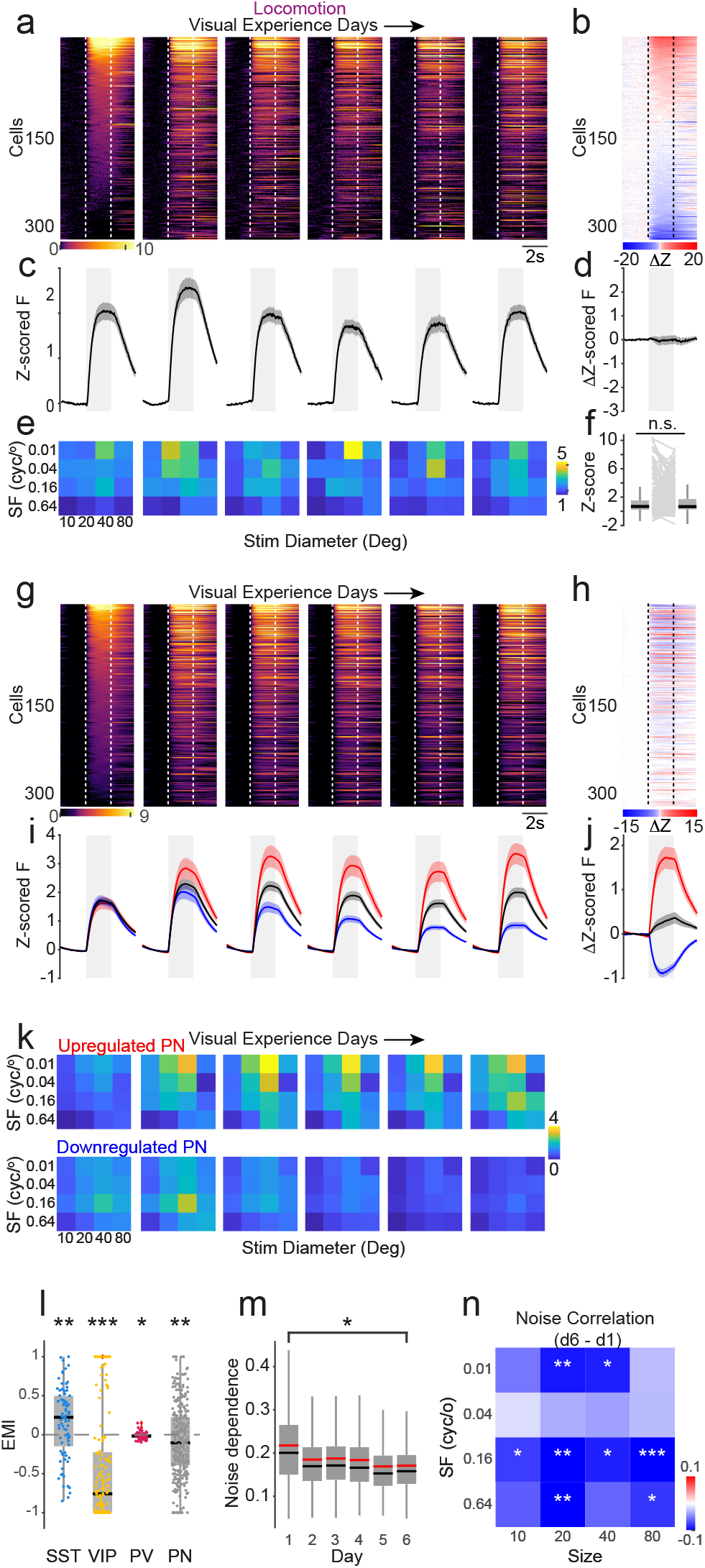
Visual exposure reshapes response properties in pyramidal neurons. (a) Raster images of average z-scored response around stimulus onset across all stimuli in Pyramidal Neurons for each day of visual exposure sorted by magnitude of response in naïve mice, restricting analysis to locomotion (n = 313 cells). (b) Raster image of the average response change around stimulus onset between naïve and visually experienced mice in Pyramidal Neurons sorted by change magnitude, restricting analysis to locomotion (n = 313 cells). (c) Average z-scored response around stimulus onset across all stimuli over days of visual exposure, restricting analysis to locomotion (n = 313 cells). (d) Average change in response between naive and visually experienced mice in Pyramidal Neurons, restricting analysis to locomotion. (e) Average z-scored response across Pyramidal Neurons for each combination of size and spatial frequency over days of visual exposure (n = 313 cells), restricting analysis to locomotion. (f) Average z-scored response across visual stimuli for each Pyramidal Neurons between naive and visually experienced mice (n = 313 cells), restricting analysis to epochs of locomotion. (g) Raster images of average z-scored response around stimulus onset across all stimuli in Pyramidal Neurons for each day of visual exposure, sorted by response amplitude on the first day (n = 313 cells). (h) Raster image of the average response change around stimulus onset across stimuli between naïve and visually experienced mice in Pyramidal Neurons, sorted by response amplitude on the first day (n = 313 cells). (i) Average z-scored response around stimulus presentation across stimuli over days of visual exposure in upregulated (red, n = 106 cells) and downregulated Pyramidal Neurons (blue, n = 148 cells) and across all Pyramidal Neurons (black, n = 313 cells). (j) Average change in response between naive and visually experienced mice in upregulated (red, n = 106 cells) and down regulated Pyramidal Neurons (blue, n = 148 cells) and across all Pyramidal Neurons (black, n = 313 cells). (k) Average z-scored response upregulated (n = 106 cells) and down regulated (n = 148 cells) Pyramidal Neurons for each combination of size and spatial frequency on the first and last days of visual exposure. (l) Experience Modulation Index (EMI) across all stimuli for SST (cyan, n = 81 cells), VIP (yellow, n = 152 cells) and PV interneurons (magenta, n = 47 cells) and for Pyramidal Neurons (gray, n = 313 cells). (m) Average noise correlation across pyramidal neurons (n = 313 cells) in each day of visual exposure. (n) Change of average noise correlation across pyramidal neurons (n = 313 cells) between the first and sixth days of visual exposure for each combination of size and spatial frequency. Data are presented as mean ± SEM; *: P<0.05, **: P<0.01, ***: P<0.001. See Supplementary Table 1 for detailed statistical analysis.

